# Structure-Based Design of a Cyclic Peptide Inhibitor of the SARS-CoV-2 Main Protease

**DOI:** 10.1101/2020.08.03.234872

**Authors:** Adam G. Kreutzer, Maj Krumberger, Chelsea Marie T. Parrocha, Michael A. Morris, Gretchen Guaglianone, James S. Nowick

## Abstract

This paper presents the design and study of a first-in-class cyclic peptide inhibitor against the SARS-CoV-2 main protease (M^pro^). The cyclic peptide inhibitor is designed to mimic the conformation of a substrate at a *C*-terminal autolytic cleavage site of M^pro^. Synthesis and evaluation of a first-generation cyclic peptide inhibitor reveals that the inhibitor is active against M^pro^ *in vitro* and is non-toxic toward human cells in culture. The initial hit described in this manuscript, UCI-1, lays the groundwork for the development of additional cyclic peptide inhibitors against M^pro^ with improved activities.

## INTRODUCTION

Antiviral drugs are desperately needed to help combat the COVID-19 pandemic caused by the Severe Acute Respiratory Syndrome coronavirus 2 (SARS-CoV-2), as well as epidemics caused by other coronaviruses in the future.^1,2^ Antiviral drugs that slow or halt viral replication can lead to a shortened time to recovery from COVID-19, offering the promise of improved mortality rates and alleviation of the tremendous strain experienced by hospitals during the COVID-19 pandemic.^3^

The main protease (M^pro^ or 3CL protease) is one of the best-characterized drug targets for coronaviruses.^4,5,6,7,8,9,10^ The SARS-CoV-2 main protease is a member of a of class homologous cysteine proteases that are needed for viral replication in diseases such as Severe Acute Respiratory Syndrome (SARS) and Middle East Respiratory Syndrome (MERS). These viruses cleave the initially translated viral polyprotein into its component proteins. Cleavage generally occurs immediately after a Gln residue, and the Gln residue is typically preceded by a hydrophobic residue, most often Leu. The residue that follows the Gln is often a small amino acid such as Ser, Ala, or Asn. M^pro^ autolytically cleaves itself from the polyprotein.^11^ Inhibiting M^pro^ activity slows or halts viral replication, offering the promise of improved clinical outcomes for COVID-19 and other coronavirus diseases. Furthermore, there are no known human proteases with similar cleavage specificity to M^pro^, suggesting that it should be possible to develop inhibitors that target M^pro^ without off-target toxicity.

The SARS-CoV-2 M^pro^ amino acid sequence is 96% identical to the SARS-CoV M^pro^ amino sequence, and the three-dimensional structure of the SARS-CoV-2 M^pro^ is highly similar to the structure of the SARS-CoV M^pro^.^12^ Peptide-based inhibitors previously developed to target the SARS-CoV M^pro^ have effectively been repurposed and modified to target the SARS-CoV-2 M^pro^ — N3 from Jin *et al*., 13b from Zhang *et al*., and 11a and 11b from Dai *et al*.^12,13,14^ These inhibitors effectively block SARS-CoV-2 replication in cell-based studies, making them promising antiviral drug candidates. While the M^pro^ inhibitors N3, 13b, 11a, and 11b have shown promise against inhibiting SARS-CoV-2 replication, additional M^pro^ inhibitors will most likely be needed for their improved properties or to be used in combination therapies.^15^

In this paper, we describe the design and preliminary evaluation of UCI-1 (**U**niversity of California, Irvine **C**oronavirus **I**nhibitor-1), a first-in-class cyclic peptide that inhibits the SARS-CoV-2 M^pro^ (Figure 1). UCI-1 is designed to mimic the conformation of a *C*-terminal autolytic cleavage site of the SARS-CoV M^pro^, a naturally occurring M^pro^ substrate. UCI-1 contains amino acid side chains from the P2, P1, P1’, and P2’ positions of the M^pro^ substrate that are designed to fill the S2, S1, S1’, and S2’ pockets of the M^pro^ active site (Figure 1A). In UCI-1, the carboxy-terminus of the P2’ residue is linked to the amino-terminus of the P2 residue with a [4-(2-aminoethyl)phenyl]-acetic acid (AEPA) group, creating a cyclophane. The (2-aminoethyl)phenyl group of AEPA is designed to act as a surrogate for a phenylalanine side chain at position P3’ and fill the S3’ pocket. Evaluation of the first generation cyclic peptide inhibitor UCI-1 in an *in vitro* M^pro^ inhibition assay reveals that UCI-1 is active against SARS-CoV-2 M^pro^ at mid-micromolar concentrations. LC/MS analysis indicates that UCI-1 resists cleavage by M^pro^, despite containing a scissile amide bond. Furthermore, UCI-1 is found to be non-toxic toward human embryonic kidney cells at concentrations that inhibit M^pro^. The following details the design, synthesis, and preliminary evaluation of UCI-1.

**Figure 1.**
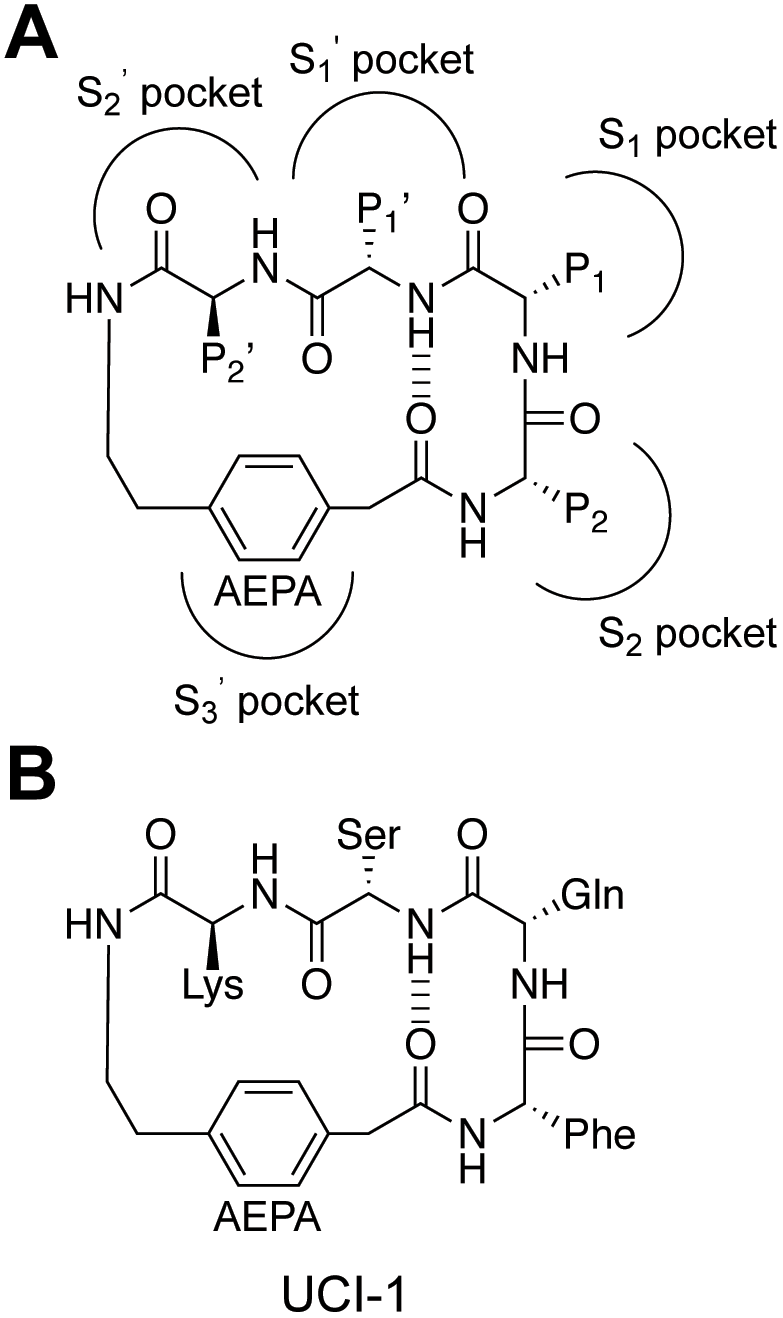
(A) Chemical structure of a general cyclic peptide inhibitor illustrating the arrangement of the P2, P1, P1’, and P2’ positions and [4-(2-aminoethyl)phenyl]-acetic acid (AEPA) and the envisioned binding interactions with the S3, S2, S1, S1’, and S2’ pockets in the M^pro^ active site. (B) Chemical structure of UCI-1.

## RESULTS AND DISCUSSION

### Design of the cyclic peptide inhibitor

We designed the cyclic peptide inhibitor based on the crystal structure of an inactive SARS-CoV M^pro^ (C145A) variant with a 10 amino-acid *C*-terminal extension corresponding to the *C*-terminal prosequence of M^pro^ (PDB 5B6O) (Figure 2).^16^ We term this M^pro^ variant “M^pro^_316_”. In the M^pro^_316_ crystal structure, *C*-terminal residues 301–310 (SGVTFQGKFK) extend into and complex with the active site of another M^pro^_316_ molecule in an adjacent asymmetric unit (Figure 2 inset). This complex reveals how the P2-P1-P1’-P2’-P3’ positions (residues 305–309, FQGKF) of the *C*-terminal autolytic cleavage site fit into the active site of M^pro^_316_.

**Figure 2.**
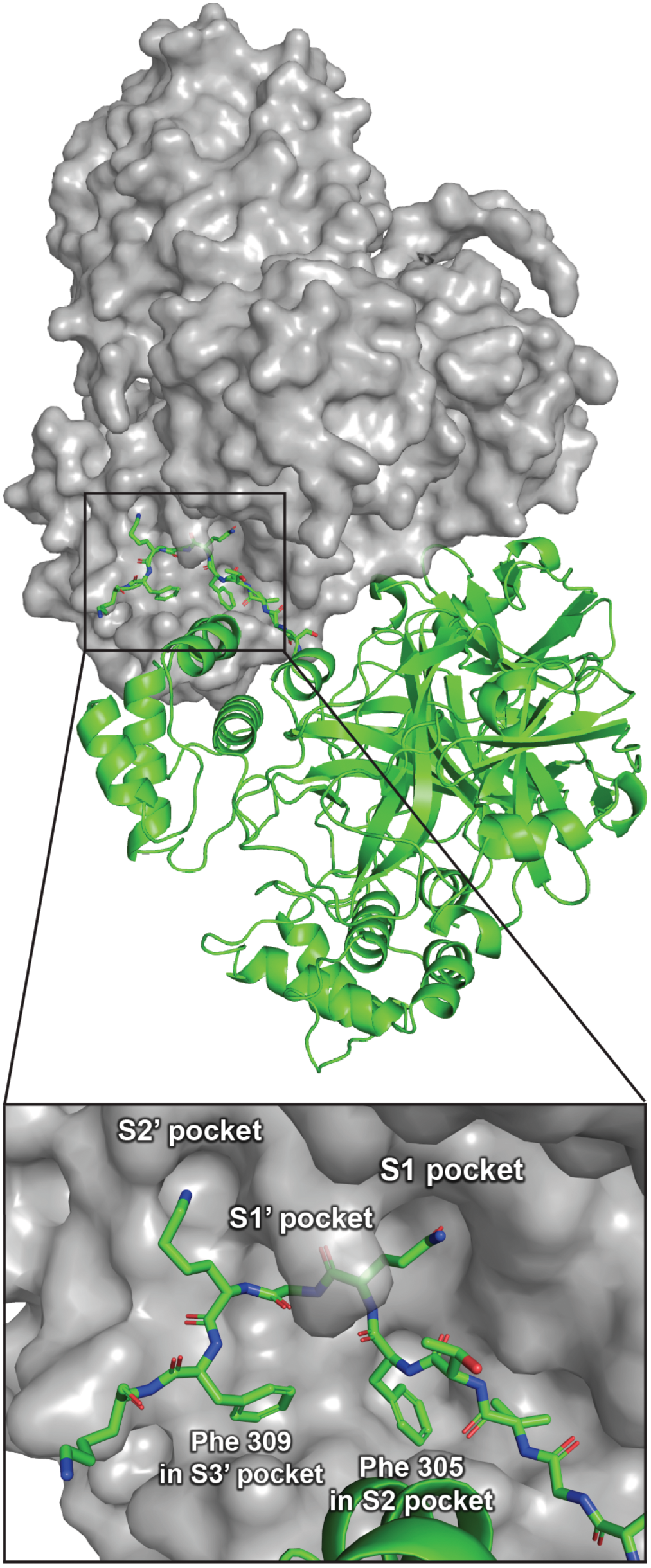
Crystal structure of M^pro^_316_ showing two M^pro^_316_ dimers in two adjacent asymmetric units (PDB 5B6O). One dimer is shown in grey surface view; the other dimer is shown in green cartoons. The inset shows a detailed view of *C*-terminal residues 301–310 of the *C*-terminal autolytic cleavage site of one M^pro^_316_ molecule in the active site of another M^pro^_316_ molecule.

The cyclic peptide inhibitor is designed to mimic the conformation that the P2-P1-P1’-P2’-P3’ residues adopt in the active site of M^pro^_316_. In the active site of M^pro^_316_, the P2-P1-P1’-P2’-P3’ residues adopt a “kinked” conformation in which the phenyl group of Phe309 at the P3’ position points toward the backbone of Phe305 at the P2 position (Figure 2 inset). To mimic this conformation, we envisioned linking the phenyl group of Phe309 to the backbone of Phe305 to create a macrocycle. To realize this design, we used the molecular visualization software PyMOL (version 2.2.2, Schrödinger) to build a model of the envisioned cyclic peptide by modifying Phe305 and Phe309 in the active site of M^pro^_316_ (Figure 3). In PyMOL, we deleted residues 301–304 to expose the amino group on Phe305; we also deleted residue 308 and the carbonyl of Phe309. We then connected the *para* position of Phe309 to the amino group of Phe305 with a CH_2_CO group to create a macrocycle. The newly created amino acid derived from Phe309 thus constitutes the amino acid AEPA.

**Figure 3.**
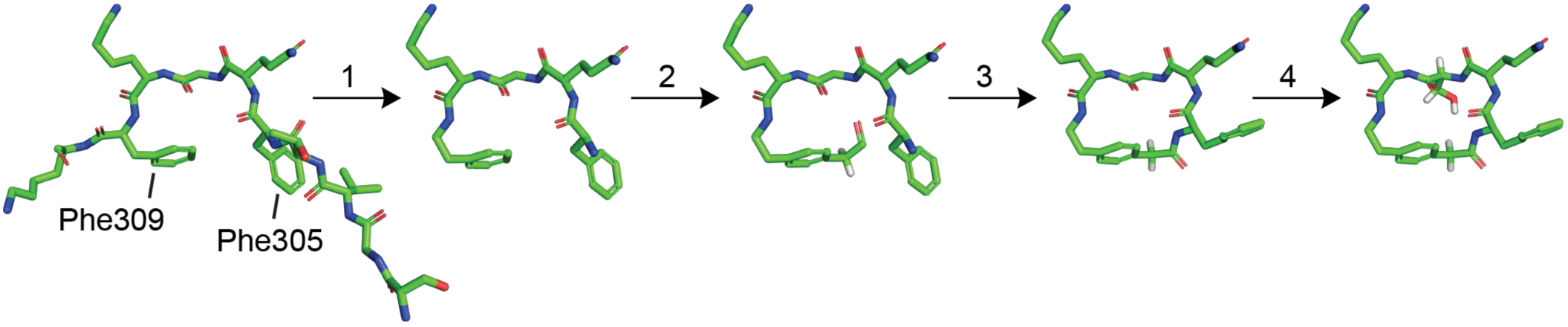
Design process for creating the cyclic peptide inhibitor UCI-1 from the *C*-terminal autolytic cleavage site that extends into the active site of the enzyme: (1) Delete residues 301– 304 and 308 as well as the carbonyl of Phe309. (2) Build a CH_2_CO group on the *para* position of the phenyl group on Phe309. (3) Create a bond between the carbonyl carbon of the newly created CH_2_CO group on Phe309 and the amino group of Phe305, and then invert the stereochemistry of Phe305. (4) Mutate Gly307 to serine.

We recognized that upon linking Phe309 and Phe305 as described above, Phe305 and Gln306 were poised to form a β-turn in which the carbonyl of AEPA hydrogen bonds with the amino group of Gly307. We envision that β-turn formation in the cyclic peptide inhibitor will promote rigidity of the cyclic scaffold. To introduce additional conformational rigidity to the macrocycle, we also mutated Gly307 to serine, which is the most common residue at the P1’ position among the 11 known SARS-CoV-2 M^pro^ cleavage sites. The resulting cyclic peptide inhibitor UCI-1 was then synthesized and further studied as described below.

### Synthesis of UCI-1

We synthesized UCI-1 by Fmoc-based solid-phase peptide synthesis of the protected linear peptide H_2_N-FQSK-AEPA-COOH on 2-chlorotrityl chloride resin followed by cleavage of the linear protected peptide from the resin and subsequent solution-phase macrocyclization and global deprotection (Scheme 1). UCI-1 was purified using reverse-phase HPLC. The synthesis and purification proceeded smoothly on a 0.1 mmol scale and yielded 22 mg of purified UCI-1 as the TFA salt. Detailed procedures for synthesis of UCI-1 are given in the Materials and Methods section.

**Scheme 1.**
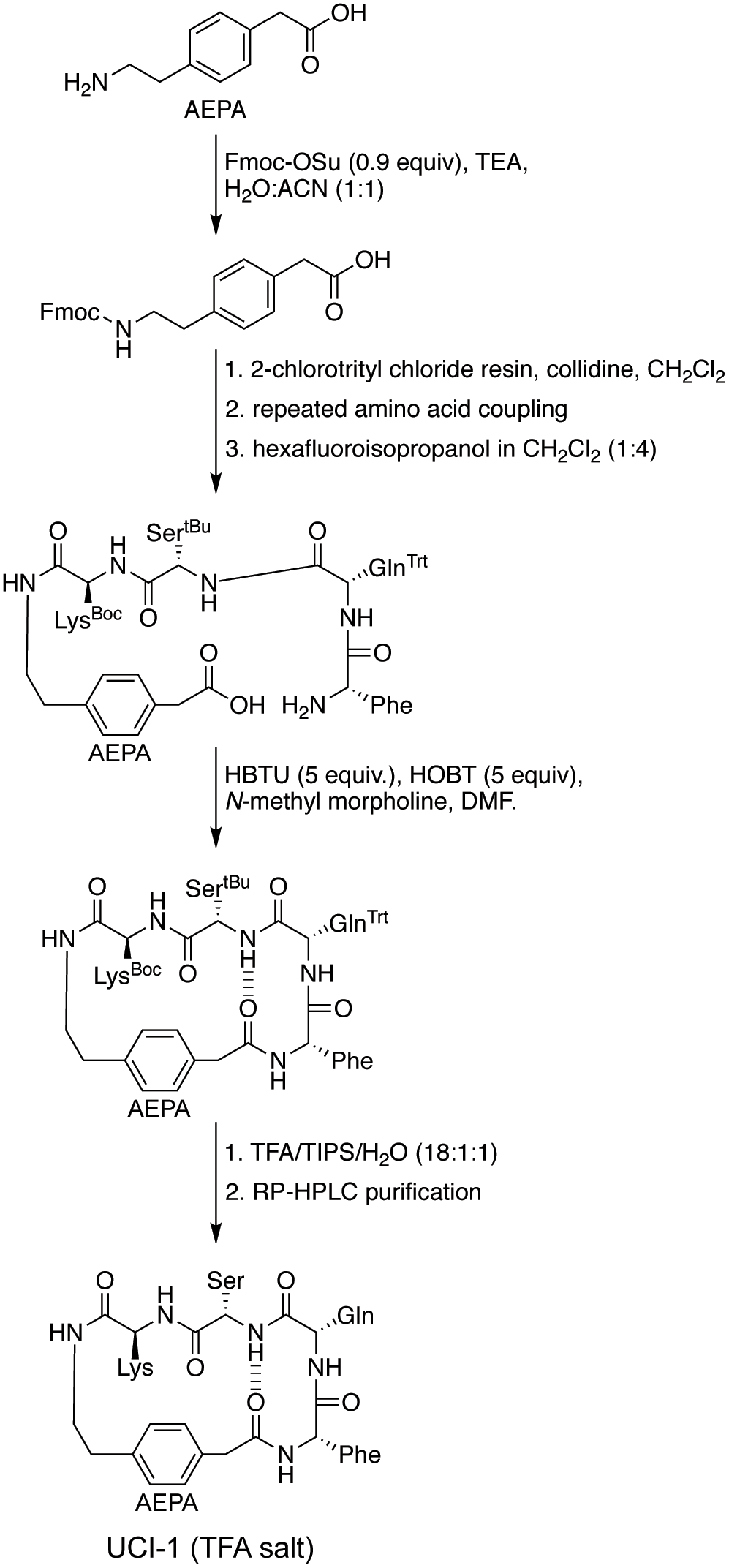
Synthesis of UCI-1.

### Enzyme inhibition assay

To evaluate the inhibitory properties of UCI-1 against M^pro^, we used an established fluorescence-based M^pro^ inhibition assay kit from BPS Bioscience. The kit includes purified SARS-CoV-2 M^pro^ as a fusion protein with maltose binding protein (MBP-M^pro^), the fluorogenic M^pro^ substrate Dabcyl-KTSAVLQSGFRKM-E(Edans)-NH_2_, and assay buffer. For the inhibition assay, we pre-incubated MBP-M^pro^ with varying concentrations of UCI-1 (32–512 µM) or with varying concentrations of the acyclic variant of UCI-1, “peptide-1a” (32–256 µM), in assay buffer for 30 min. We then added the fluorogenic M^pro^ substrate and monitored MBP-M^pro^ activity in a continuous kinetic assay using wavelengths of 360 nm and 460 nm for excitation and emission. In each well, the concentration of MBP-M^pro^ was 0.2 µM and the concentration of the fluorogenic substrate was 50 µM. Figure S1 shows the results of the results of the continuous kinetic assay of UCI-1.

Initial rates for MBP-M^pro^ activity in the presence or absence of UCI-1 or peptide-1a were obtained by fitting the linear portions of the curves from the continuous kinetic assay to a straight line. UCI-1 is active against MBP-M^pro^ with an IC_50_ value of ∼150 µM (Figure 4A). In contrast, the linear control peptide-1a shows little or no inhibition at concentrations at or below 256 µM (Figure 4B). There appears to be a slight reduction in rate of cleavage of the fluorogenic substrate upon addition of 256 µM peptide-1a. This reduction in rate may reflect either slight inhibition of MBP-M^pro^ or that the linear peptide acts as a competitive substrate at high concentrations. These findings indicate that the cyclic structure of UCI-1 is critical for its activity.

**Figure 4.**
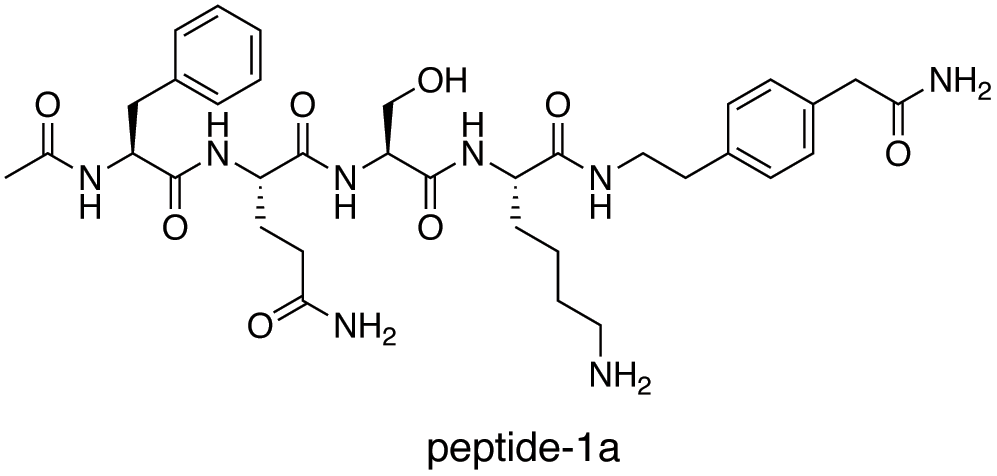
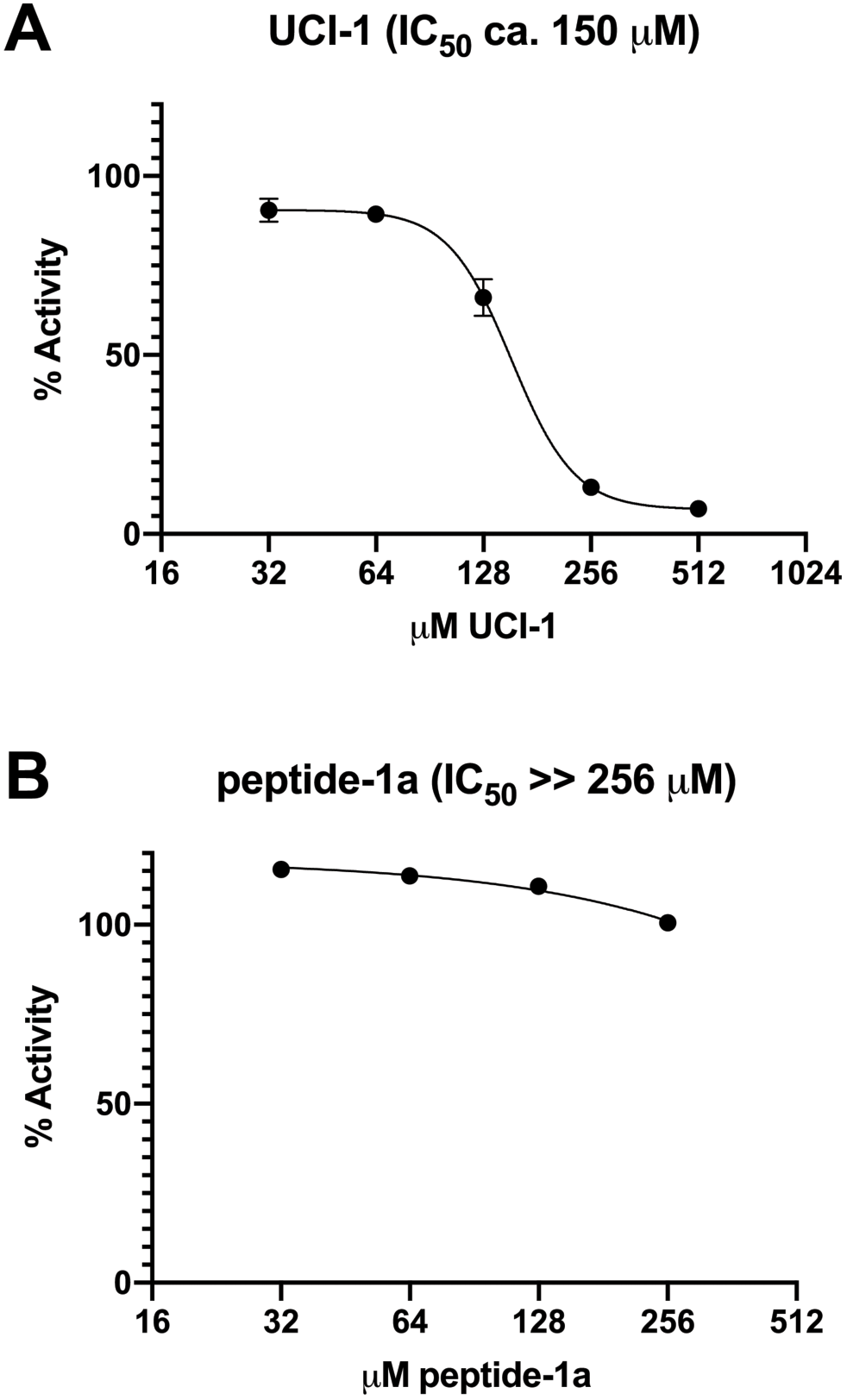
Enzyme inhibition assays. The activity of MBP-M^pro^ was measured in the presence of increasing concentrations of UCI-1 (A) or lin-UCI-1 (B). Dose response curves for IC_50_ values were determined by non-linear regression. All data are shown as mean ± s.d., *n* = 3 replicates.

### Assessment of cleavage of UCI-1 by M^pro^

The amide bond between residues at positions P1 and P1’ of UCI-1 has the potential to be cleaved by M^pro^ because residues at these positions correspond to the cleavage site of M^pro^ substrates. To determine whether M^pro^ cleaves UCI-1, we used LC/MS to analyze UCI-1 in the well solution from the 96-well plate of the enzyme inhibition assay after 24 hours. To aide in LC/MS identification of the cleavage product of UCI-1, we synthesized the authentic UCI-1 M^pro^ cleavage product “peptide-1b” and spiked the well solution with peptide-1b.

UCI-1 does not appear to be cleaved by M^pro^ (Figure 5). LC/MS analysis of the enzyme inhibition assay well solution with 16 µM UCI-1 spiked with 0.1 µM peptide-1b shows a small peak at 2.09 minutes that corresponds to peptide-1b (Figure 5B). The peak at 2.09 minutes is absent in the LC/MS trace acquired before spiking the well solution with peptide-1b (Figure 5A), indicating that there is not appreciable cleavage of UCI-1 by M^pro^. The ion current at 2.09 minutes in the LC/MS trace acquired before spiking with peptide-1b shows no evidence of the UCI-1 cleavage product (Figure 5C), whereas the ion current at 2.09 minutes in the LC/MS trace acquired after spiking with peptide-1b shows the mass of peptide-1b (Figure 5D), providing additional evidence that UCI-1 resists cleavage by M^pro^. These findings confirm that UCI-1 acts as an inhibitor and not a competitive substrate. The resistance of UCI-1 to cleavage by M^pro^ is consistent with the observations by others that cyclic peptides often resist proteolytic cleavage.^17,18,19,20,21,22^

**Figure 5.**
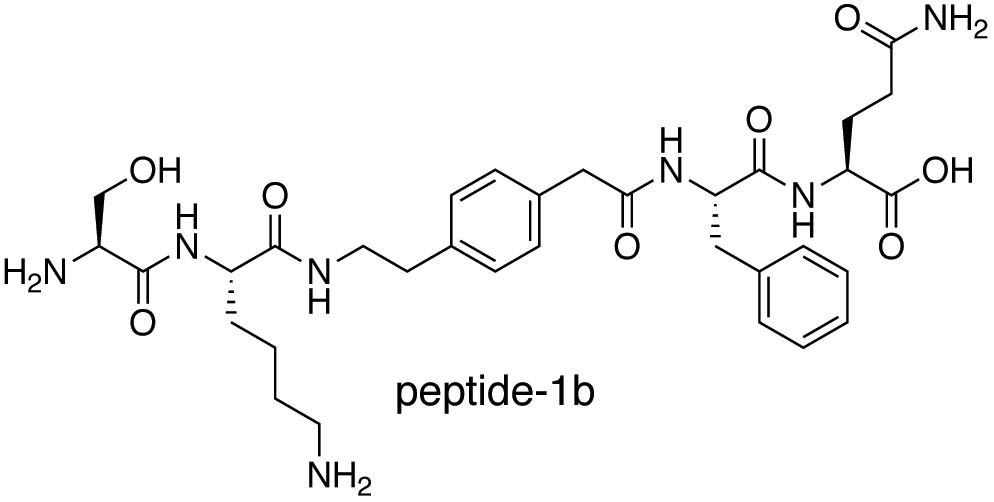
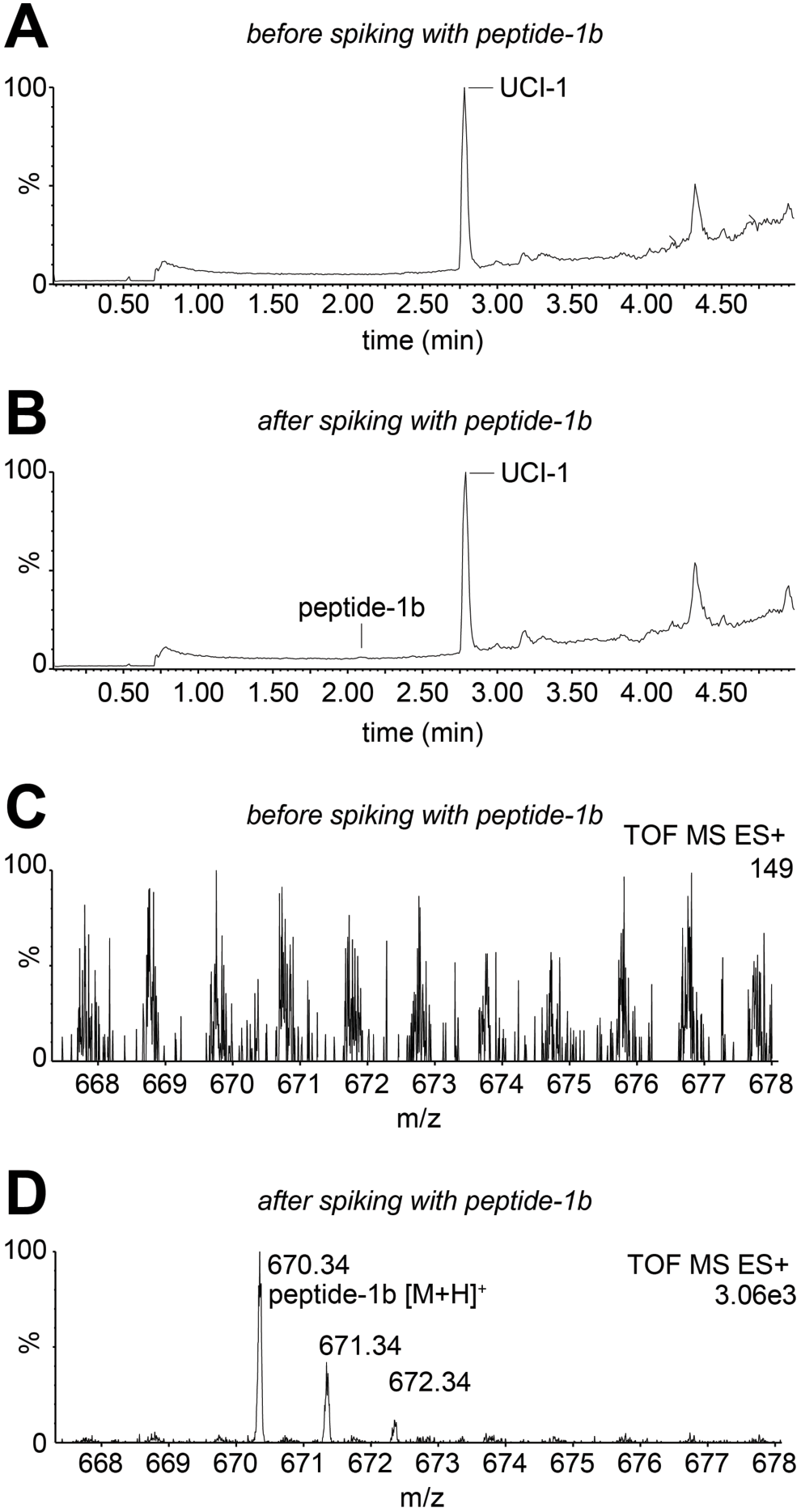
Assessment of cleavage of UCI-1 by M^pro^ using LC/MS. (A) LC/MS total ion current spectrum of well solution from M^pro^ inhibition assay with 16 µM UCI-1. (B) LC/MS total ion current spectrum of well solution from M^pro^ inhibition assay with 16 µM UCI-1 spiked with 0.1 µM peptide-1b. (C) Ion current for peptide-1b (2.09 min) in LC/MS spectrum of well solution from M^pro^ inhibition assay with 16 µM UCI-1. (D) Ion current for peptide-1b (2.09 min) in LC/MS spectrum of well solution from M^pro^ inhibition assay with 16 µM UCI-1 spiked with 0.1 µM peptide-1b.

### Cytotoxicity of UCI-1

To evaluate whether UCI-1 is cytotoxic, we exposed human embryonic kidney (HEK-293) cells to varying concentrations of UCI-1 (0–256 µM) for 72 hours, and then assessed cell death using a lactase dehydrogenase (LDH) assay. At the highest concentration evaluated (256 µM), UCI-1 elicits little to no LDH release from HEK-293 cells, indicating that UCI-1 is not cytotoxic at concentrations up to 256 µM (Figure 6).

**Figure 6.**
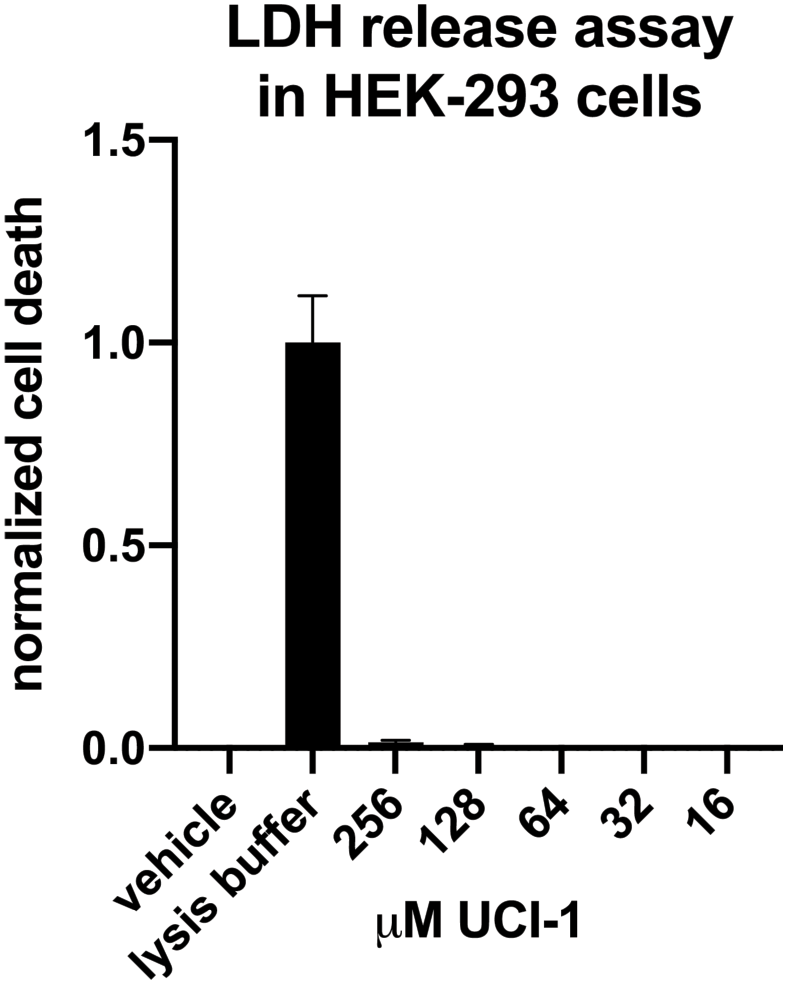
LDH release assay of UCI-1 in HEK-293 cells. All data are shown as mean ± s.d., *n* = 5 replicates. Deionized water (vehicle) was used as a negative control.

### Conformational analysis and docking of UCI-1

To better understand the three-dimensional structure of UCI-1 and how it interacts with the active site of the SARS-CoV-2 M^pro^, we performed conformational analysis and docking studies. Conformational searching of UCI-1 (MacroModel with the MMFFs force field and GB/SA water) revealed that UCI-1 adopts a global minimum energy conformation that resembles the kinked conformation that residues 305–309 adopt in the active site of M^pro^_316_ (Figures 2 and 7A). In the global minimum energy conformation, the AEPA residue acts as a rigid spacer, with the Phe, Gln, Ser, and Lys forming a bridge. As we had envisioned, the Phe and Gln residues adopt a β-turn conformation, with Phe at the i+1 position and Gln at the i+2 position. The Phe side chain is well situated to fit in the S2 pocket, and the Gln side chain is well situated to fit in the S1 pocket. The Ser, Lys, and AEPA residues, in turn, are poised to occupy the S1’, S2’, and S3’ pockets. The macrocyclic scaffold of the cyclic peptide inhibitor appears to be particularly rigid. In the conformational search, the peptide backbones of most of the conformers adopt the conformation described above, differing only in side chain geometry and the type of β-turn formed by Phe305 and Gln306 (Figure 7B). Docking of the lowest energy conformer of UCI-1 with the SARS-CoV-2 M^pro^ (PDB 6YB7) using Autodock Vina reveals that the inhibitor binds the active site of M^pro^ in the envisioned manner (Figure 7C).^23^

**Figure 7.**
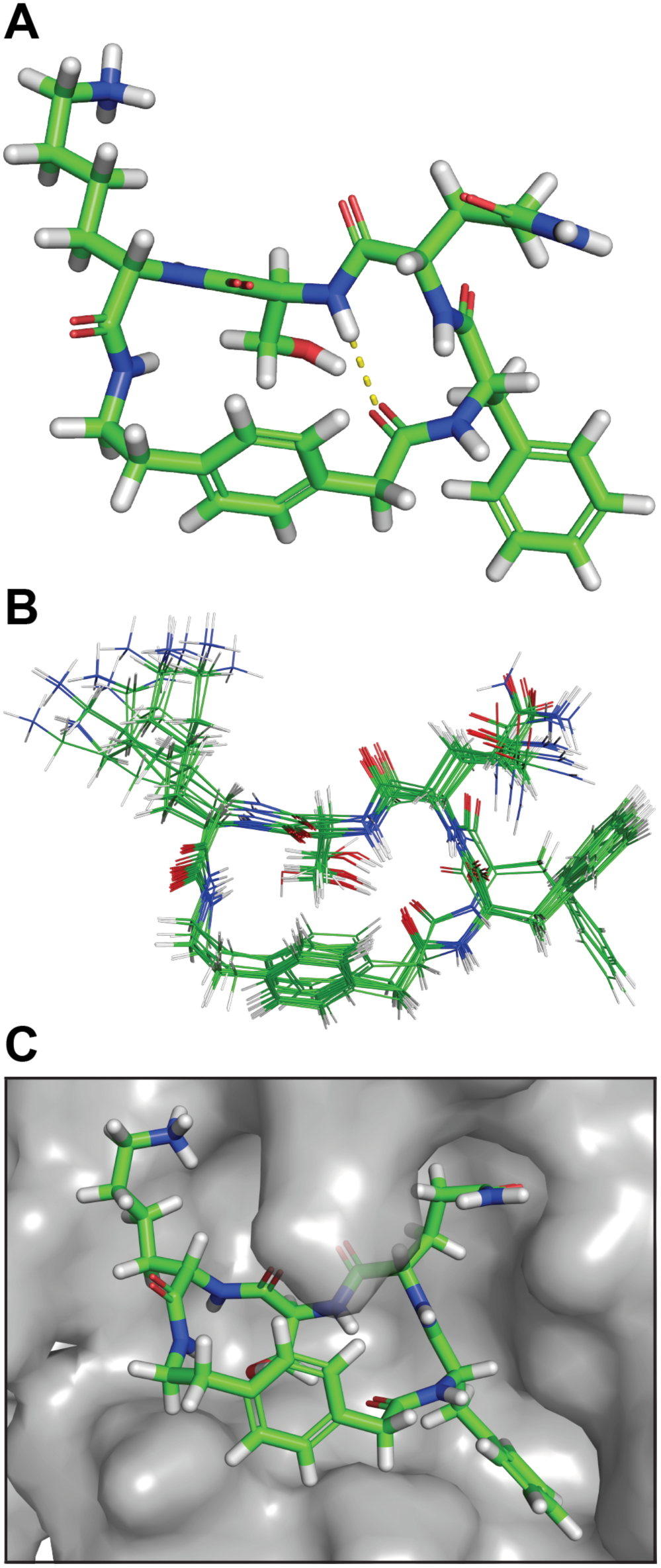
(A) Lowest energy conformer of UCI-1. (B) Superposition of the 20 lowest energy conformers from conformational searching. The difference in energy between the lowest and highest energy conformers among these 20 is 7.0 kJ/mol. (C) UCI-1 in complex with the SARS-CoV-2 M^pro^ active site generated in Autodock Vina. The SARS-CoV-2 M^pro^ crystal structure with PDB accession number 6YB7 was used in the docking study.

## CONCLUSIONS

The design and preliminary evaluation of UCI-1 demonstrates that cyclic peptides that mimic the conformation of linear peptide substrates of M^pro^ can be developed as inhibitors against M^pro^. Almost all of the M^pro^ inhibitors that have been reported are linear peptides that contain a “warhead” that forms a covalent bond with the active cysteine of M^pro^. While the activity of UCI-1 is modest compared to other known M^pro^ inhibitors, the design and preliminary evaluation of UCI-1 lays the groundwork for developing additional cyclic peptide inhibitor analogs of UCI-1 with improved activity against M^pro^. Design and development of next generation UCI-1 analogs will likely produce better inhibitors. Due to the urgency of COVID-19, we elected to share our initial hit, UCI-1, in this preprint. We hope that doing so will encourage others to also begin thinking about cyclic peptide inhibitors as promising drug candidates. We are currently pursuing next generation analogs of UCI-1 predicted to exhibit enhanced activity against M^pro^, and will report our findings from these pursuits in due course.

## MATERIALS AND METHODS

### Synthesis of 2-(4-(2-((((9H-fluoren-9-yl)methoxy)carbonyl)amino)ethyl)phenyl)acetic acid (Fmoc-AEPA)

A 50 mL round-bottom flask equipped with a magnetic stirring bar was charged with 100 mg (0.55 mmol, 1 equiv) of 2-(4-(2-aminoethyl)phenyl)acetic acid dissolved in 10 mL H_2_O. 0.156 mL (1.10 mmol, 2 equiv) of Et_3_N was added. 160 mg of Fmoc-OSu (0.50 mmol, 0.9 equiv) was dissolved in 10 mL CH_3_CN and added to the reaction mixture. The reaction was run for 2 hours at room temperature. While it was running, the reaction was monitored by TLC (3:1 EtOAc/hexanes + 10 % MeOH, R_f_ = 0.44) to determine the consumption of starting material (R_f_ = 0.77) and an appearance of fulvene (R_f_ = 0.81). 10 mL of EtOAc was then added to the reaction mixture and the organic layer was removed. The aqueous layer was acidified with 30 mL 1 M HCl, and then 10 mL of EtOAc was added. The organic layer was washed with water and brine, dried over MgSO_4_, and solvent was evaporated *in vacuo* to afford a white powder (70 %, 0.140 g). The Fmoc-AEPA was used in solid-phase peptide synthesis without further purification. The product contains a minor contaminant (< 10 %) of Fmoc-AEPA-AEPA-OH, as detected by ^1^H NMR spectroscopy. HRMS (ESI-TOF) *m/z*: [M+Na]^+^ calcd for C_25_H_23_NO_4_ 424.1525 found 424.1507.

### Synthesis of UCI-1

2-Chlorotrityl chloride resin (200 mg, 1.6 mmol/g) was added to a Bio-Rad Poly-Prep chromatography column (10 mL). The resin was suspended in dry CH_2_Cl_2_ (10 mL) and allowed to swell for 30 min. The solution was drained from the resin and a solution of Fmoc-AEPA-OH (0.50 equiv, 64 mg, 0.16 mmol) in 6% (v/v) 2,4,6-collidine in dry CH_2_Cl_2_ (8 mL) was added immediately and the suspension was gently agitated for 12 h. The solution was then drained and a mixture of CH_2_Cl_2_/MeOH/*N,N*-diisopropylethylamine (DIPEA) (17:2:1, 10 mL) was added immediately. The mixture was gently agitated for 1 h to cap the unreacted 2-chlorotrityl chloride resin sites. The resin was then washed with dry CH_2_Cl_2_ (2x) and dried by passing nitrogen through the vessel. This procedure yielded 0.26 mmol/g of loaded resin.

The Fmoc-AEPA-2-chlorotrityl resin was transferred to a peptide synthesis coupling vessel and subjected to cycles of peptide coupling with Fmoc-protected amino acid building blocks. The linear peptide was synthesized from the *C*-terminus to the *N*-terminus. Each coupling cycle consisted of i. Fmoc-deprotection with 20% (v/v) piperidine in DMF for 5 min (2x), ii. washing with DMF (3x), iii. coupling of the amino acid (5 equiv) in the presence of HCTU (4.5 equiv) and 20% (v/v) *N*-methylmorpholine (2,4,6-collidine) in DMF for 10–20 min. iv. washing with DMF (3x). After coupling of the last amino acid, the terminal Fmoc group was removed with 20% (v/v) piperidine in DMF. The resin was transferred from the coupling vessel to a Bio-Rad Poly-Prep chromatography column.

The linear peptide was cleaved from the resin by agitating the resin for 1 h with a solution of 1,1,1,3,3,3-hexafluoroisopropanol (HFIP) in CH_2_Cl_2_ (1:4, 7 mL).^24^ The suspension was filtered and the filtrate was collected in a 250-mL round-bottomed flask. The resin was washed with additional HFIP in CH_2_Cl_2_ (1:4, 7 mL) and then with CH_2_Cl_2_ (2×10 mL). The combined filtrates were concentrated by rotary evaporation to give a white solid. The white solid was further dried by vacuum pump to afford the crude protected linear peptide, which was cyclized without further purification.

The crude protected linear peptide was dissolved in dry DMF (150 mL). HOBt (5 equiv) and HBTU (5 equiv) were added to the solution. NMM (12 equiv) was added to the solution and the mixture was stirred under nitrogen for 24 h. The mixture was then concentrated under reduced pressure to afford the crude protected cyclic peptide.

The protected cyclic peptide was dissolved in TFA/triisopropylsilane (TIPS)/H_2_O (18:1:1, 20 mL) in a 250-mL round-bottomed flask equipped with a nitrogen-inlet adaptor. The solution was stirred for 1.5 h. The reaction mixture was then concentrated by rotary evaporation under reduced pressure to afford the crude cyclic peptide as a thin yellow film on the side of the round-bottomed flask. The crude cyclic peptide was immediately subjected to purification by reverse-phase HPLC (RP-HPLC).

The peptide was dissolved in H_2_O and acetonitrile (7:3, 10 mL), and the solution was filtered through a 0.2 µm syringe filter and purified by RP-HPLC (gradient elution with 10–30% CH_3_CN over 50 min). Pure fractions were concentrated by rotary evaporation and lyophilized. The synthesis of UCI-1 yielded 22 mg of the peptide as the TFA salt.

### Synthesis of peptide-1a

Rink amide AM resin (300 mg, 0.68 mmol/g) was added to a peptide synthesis coupling vessel. The resin was suspended in dry DMF (10 mL) and allowed to swell for 30 min. The solution was drained from the resin and the Fmoc group was removed with 20% (v/v) piperidine in DMF for 20 min. The solution was drained and washed with DMF (5x). Fmoc-AEPA-OH (0.50 equiv, 40.8 mg, 0.102 mmol), HATU (0.5 equiv), and HOAt (0.5 equiv) in 20% (v/v) 2,4,6-collidine in dry DMF (8 mL) was then added to the resin mixed for 12 h. The solution was then drained, washed with DMF (3x) and a mixture of acetic anhydride/pyridine (3:2, 10 mL) was added. The resin was mixed for 15 min to cap the unreacted resin sites. The resin was then washed with DMF (3x). This procedure yielded 0.17 mmol/g of loaded resin.

The Fmoc-AEPA-Rink amide AM resin was subjected to cycles of peptide coupling with Fmoc-protected amino acid building blocks as described above, ending with cleavage of the terminal Fmoc. The resin was then washed with CH_2_Cl_2_ (3x) and then dried by pushing N_2_ gas through the Poly-Prep column. The peptide was cleaved from the resin and globally deprotected by mixing the dried resin with TFA/triisopropylsilane (TIPS)/H_2_O (18:1:1, 10 mL) and gently rocking for 2.5 hours. The peptide was drained into a glass beaker, precipitated in cold ether, and subjected to purification by RP-HPLC as described above. The synthesis of peptide-1a yielded 17 mg of the peptide as the TFA salt.

### Synthesis of peptide-1b

2-Chlorotrityl chloride resin (300 mg, 1.6 mmol/g) was added to a Bio-Rad Poly-Prep chromatography column (10 mL). The resin was suspended in dry CH_2_Cl_2_ (10 mL) and allowed to swell for 30 min. The solution was drained from the resin and a solution of Fmoc-Gln(Trt)-OH (0.50 equiv, 146.57 mg, 0.24 mmol) in 6% (v/v) 2,4,6-collidine in dry CH_2_Cl_2_ (8 mL) was added immediately and the suspension was gently agitated for 12 h. The solution was then drained and a mixture of CH_2_Cl_2_/MeOH/*N,N*-diisopropylethylamine (DIPEA) (17:2:1, 10 mL) was added immediately. The mixture was gently agitated for 1 h to cap the unreacted 2-chlorotrityl chloride resin sites. The resin was then washed with dry CH_2_Cl_2_ (2x) and dried by passing nitrogen through the vessel. This procedure yielded 0.36 mmol/g of loaded resin.

The Fmoc-Gln(Trt)-2-chlorotrityl resin was subjected to cycles of peptide coupling with Fmoc-protected amino acid building blocks as described above, ending with cleavage of the terminal Fmoc. The resin was then washed with CH_2_Cl_2_ (3x) and then dried by pushing N_2_ gas through the Poly-Prep column. The peptide was cleaved from the resin and globally deprotected by mixing the dried resin with TFA/triisopropylsilane (TIPS)/H_2_O (18:1:1, 10 mL) and gently rocking for 2.5 hours. The peptide was drained into a glass beaker, precipitated in cold ether, and subjected to purification by RP-HPLC as described above. The synthesis of peptide-1b yielded 13 mg of the peptide as the TFA salt.

### Enzyme inhibition assay

A proprietary buffer containing detergent from BPS Bioscience was used for the inhibition assays. The substrate with the cleavage sites of M^pro^ (indicated by the arrow, ↓), Dabcyl-KTSAVLQ↓SGFRKM-E(Edans)-NH_2_ and (BPS Bioscience), was used in the fluorescence resonance energy transfer (FRET)-based continuous kinetic assay, using a back-walled 96-well plate. The dequenching of the Edans fluorescence due to the cleavage of the substrate by MBP-M^pro^ was monitored at 460 nm with excitation at 360 nm, using a Varioskan LUX fluorescence spectrophotometer (Thermo Fisher) using the top-read mode. Stock solutions (10 mg/mL) of UCI-1 and peptide-1a were prepared gravimetrically by dissolving the peptides in deionized H_2_O. For the determination of the IC_50_, 0.2 µM SARS-CoV-2 MBP-M^pro^ was incubated with UCI-1 or peptide-1a at various concentrations (32–512 µM for UCI-1 and 32–256 µM peptide-1a) in assay buffer at room temperature for 30 min. Afterward, the reaction was initiated by adding the FRET peptide substrate at a 50 µM final concentration (final volume: 50 µL). The IC_50_ value for UCI-1 was determined using the GraphPad Prism 8.4.3 software (GraphPad) by plotting the initial rates. Measurements of enzymatic activity were performed in triplicate and are presented as the mean ± standard deviations (s.d.).

### LC/MS

LC/MS analysis of the pooled 16 µM UCI-1 well solutions from the enzyme inhibition assay was performed on a Waters Xevo Qtof G2XS equipped with a C4 column. For both LC/MS traces, 10 µL of the well solution (diluted 1:100 in deionized H_2_O) was injected on the column. For the spiking experiment, a 10 mg/mL stock solution of peptide-1b was prepared gravimetrically in deionized H_2_O, and an aliquot of the stock solution was diluted in deionized H_2_O to create a 10 µM working solution. A 1-µL aliquot of the 10 µM working solution of peptide-1b was added to 100 µL of the well solution to achieve a final peptide-1b concentration of 0.1 µM.

### LDH release assay

The LDH release assay was performed using the Pierce LDH Cytotoxicity Assay Kit from Thermo Scientific. Experiments were performed in replicates of five, and an additional 10 wells were used for controls. Cells were cultured in the inner 60 wells (rows B–G, columns 2–11) of the 96-well plate. DMEM:F12 media (100 µL) was added to the outer wells (rows A and H and columns 1 and 12), in order to ensure the greatest reproducibility of data generated from the inner wells. A 10-mg/mL stock solution of UCI-1 was prepared gravimetrically in sterile deionized H_2_O that had been passed through a 0.2 µm nylon syringe filter. The stock solution was used to create a 2560 µM solution of UCI-1, which was serially diluted in sterile deionized H_2_O to create 10X working solutions of UCI-1.

HEK-293 cells were plated in a 96-well plate at 15,000 cells per well. Cells were incubated in 100 µL of a 1:1 mixture of DMEM:F12 media supplemented with 10% fetal bovine serum, 100 U/mL penicillin, and 100 µg/mL streptomycin at 37 °C in a 5% CO_2_ atmosphere and allowed to adhere to the bottom of the plate for 24 hours. After 24 hours, the culture media was removed and replaced with 90 µL of serum-free DMEM:F12 media. A 10-µL aliquot of the working solutions of UCI-1 was added to each well, for well concentrations of 256 µM to 32 µM. Experiments were run in replicates of five. Five wells were used as controls and received 10-µL aliquots of sterile deionized water (vehicle). Another five wells were left untreated, to be subsequently used as controls with lysis buffer for the LDH release assay. Cells were incubated at 37 °C in a 5% CO_2_ atmosphere for 72 hours.

After 72 hours, 10 µL of 10x lysis buffer — included with the assay kit — was added to the five untreated wells, and the cells were incubated for an additional 45 min. After 45 min, a 50-µL aliquot of the supernatant media from each well was transferred to a new 96-well plate and 50 µL of LDH substrate solution, prepared according to manufacturer’s protocol, was added to each well. The treated plates were stored in the dark for 30 min. The absorbance of each well was measured at 490 and 680 nm (A_490_ and A_680_). Data were processed by calculating the differential absorbance for each well (A_490_−A_680_) and comparing those values to those of the lysis buffer controls and the untreated controls:

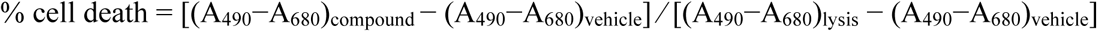

### Docking of UCI-1 to SARS-CoV-2 M^pro^

The model of the SARS-CoV-2 M^pro^ was generated as follows: Starting coordinates of SARS-CoV-2 M^pro^ were generated from the SARS-CoV-2 M^pro^ crystallographic structure (PDB 6YB7) using PyMOL and saved as a new PDB file. In PyMOL the dimethyl sulfoxide molecule that sits in the active site of 6YB7 was deleted. iBabel was used to convert the UCI-1 minimized structured PDB file into a PDBQT file prior to docking. Docking was performed using AutoDock Tools and AutoDock Vina. In AutoDock Tools, a grid was chosen to encompass the active site of SARS-CoV-2 M^pro^ in the size of 25×25×25 Å and with the coordinates x = 9.250, y = −5.944, z = 18.944. SARS CoV-2 M^pro^ was treated as a rigid receptor in these calculations. The lowest energy cluster, as determined by AutoDock Vina was chosen to represent the SARS-CoV-2 M^pro^ UCI-1 interaction model in Figure 7C.

## AUTHOR CONTRIBUTIONS

A.G.K. and J.S.N. conceived and designed UCI-1 and the research. A.G.K., M.K., and M.A.M. synthesized Fmoc-AEPA-OH. A.G.K., C.M.T.P., and M.A.M. synthesized UCI-1, peptide-1a, and peptide-1b. A.G.K. performed the enzyme inhibition assays and the LC/MS experiments. G.G. performed the LDH release assays. A.G.K. and J.S.N. performed the conformational analysis. M.K. performed the docking study. A.G.K. and J.S.N. wrote the manuscript. All authors read and approved the manuscript.

## AUTHOR INFORMATION

### Notes

The authors declare no competing financial interest.

## ADDITIONAL INFORMATION

Supplementary information accompanies this paper.

## Notes

### Competing Interest Statement

The authors have declared no competing interest.

### Summary of Updates

There is a mistake in the original submission. In the subsection "Conformational analysis and docking of UCI-1", "D-Phe" should instead be called "Phe".

