## Supplementary Information for "Structure-Based Design of a Cyclic Peptide Inhibitor of the SARS-CoV-2 Main Protease"

**This PDF includes:**

**Supplementary Figures**

Figure S1. Continuous kinetic assay of UCI-1 inhibition of the SARS-CoV-2 –MBP $M^{pro}$. S2

**Supplementary Procedures** S2

Synthesis and characterization of Fmoc-AEPA-OH. S2

**Peptide Characterization Data**

Characterization of UCI-1. S12

Characterization of peptide-1a. S14

Characterization of peptide-1b. S16

### Supplementary Figures

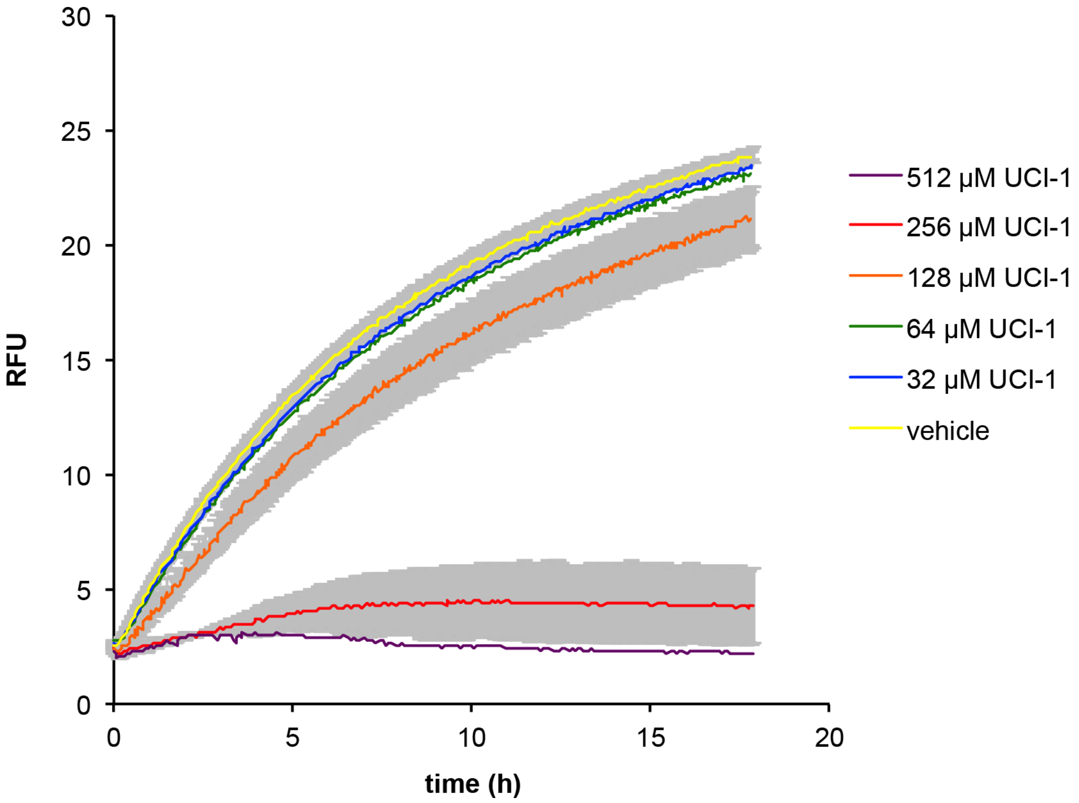


**Figure S1.** Continuous kinetic inhibition assay of UCI-1 against MBP-$M^{pro}$. For clarity, error bars (± s.d.) are only shown for 256 μM UCI-1, 128 μM UCI-1, and vehicle.

### Supplementary Procedures

#### Synthesis of 2-(4-(2-((((9H-fluoren-9-yl)methoxy)carbonyl)amino)ethyl)phenyl)acetic acid (Fmoc-AEPA)

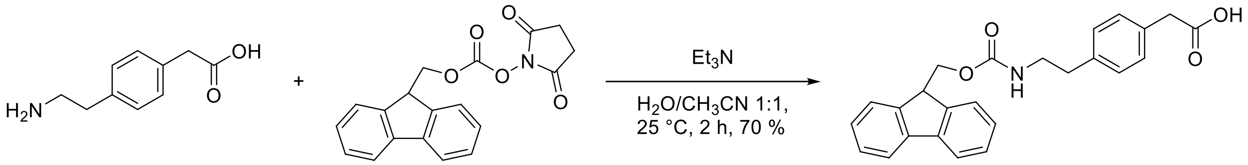


A 50 mL round-bottom flask equipped with a magnetic stirring bar was charged with 100 mg (0.55 mmol, 1 equiv) of 2-(4-(2-aminoethyl)phenyl)acetic acid dissolved in 10 mL $H_2O$. 0.156 mL (1.10 mmol, 2 equiv) of $Et_3N$ was added. 160 mg of Fmoc-OSu (0.50 mmol, 0.9 equiv) was dissolved in 10 mL $CH_3CN$ and added to the reaction mixture. The reaction was run for 2 hours at room temperature. While it was running, the reaction was monitored by TLC (3:1

EtOAc/hexanes + 10 % MeOH, $R_f$ = 0.44) to determine the consumption of starting material and ($R_f$ = 0.77) and an appearance of fulvene ($R_f$ = 0.81). 10 mL of EtOAc was then added to the reaction mixture and the organic layer was removed. The aqueous layer was acidified with 30 mL 1 M HCl, and then 10 mL of EtOAc was added. The organic layer was washed with water and brine, dried over $MgSO_4$, and solvent was evaporated *in vacuo* to afford a white powder (70 %, 0.140 g). The Fmoc-AEPA was used in solid-phase peptide synthesis without further purification. The product contains a minor contaminant (< 10 %) of Fmoc-AEPA-AEPA-OH, as detected by $^1$H NMR spectroscopy. HRMS (ESI-TOF) *m/z*: $[M+Na]^+$ calcd for $C_{25}H_{23}NO_4$ 424.1525 found 424.1507.

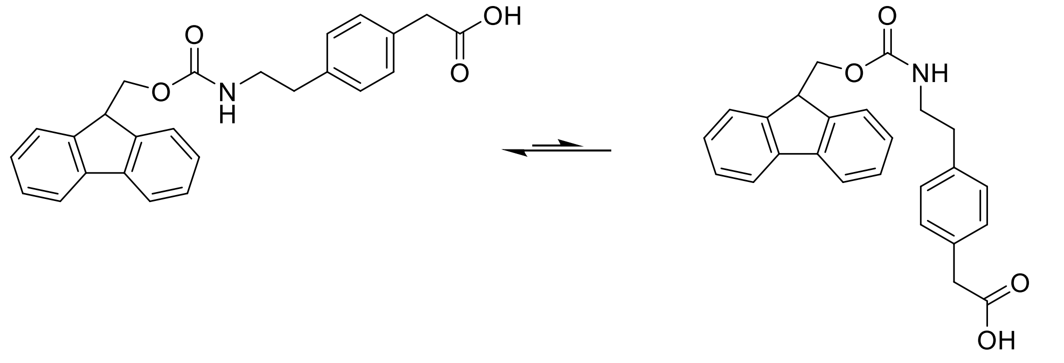


Rotamer 1: Rotamer 2 = ~ 5:1

Rotamer 1: **$^1$H NMR (600MHz, DMSO-*d6*)**: δ 12.17 (s, 1H), 7.89 (d, J = 7.6 Hz, 2H), 7.68 (d, J = 7.3 Hz, 2H), 7.41 (t, J = 7.1 Hz, 2H), 7.38 (t, J = 5.3 Hz, 1H), 7.33 (t, J = 7.3 Hz, 2H), 4.30 (d, J = 3.2 Hz, 2H), 4.20 (t, J = 6.6 Hz, 1H), 3.51 (s, 2H), 3.19 (q, J = 6.3 Hz, 2H), 2.69 (t, J = 7.1 Hz, 2H). **$^{13}$C NMR (150 MHz, DMSO-*d6*)**: δ 173.3, 144.4, 141.3, 138.1, 133.2, 129.8, 129.0, 128.1, 127.5, 125.7, 120.6, 65.7, 47.3, 42.3, 40.8, 40.5, 35.4.

Rotamer 2: **$^1$H NMR (600MHz, DMSO-*d6*)**: δ 12.17 (s, 1H), 7.89 (d, J = 7.6 Hz, 2H), 7.68 (d, J = 7.3 Hz, 2H), 7.41 (t, J = 7.1 Hz, 2H), 7.33 (t, J = 7.3 Hz, 2H), 6.84 (t, J = 5.6 Hz, 1H), 4.30 (d, J = 3.2 Hz, 2H), 4.20 (t, J = 6.6 Hz, 1H), 3.51 (s, 2H), 2.92 (q, J = 6.3 ppm, 2H), 2.36 (t, J = 7.1 Hz, 2H).

**$^1$H NMR assignment:**

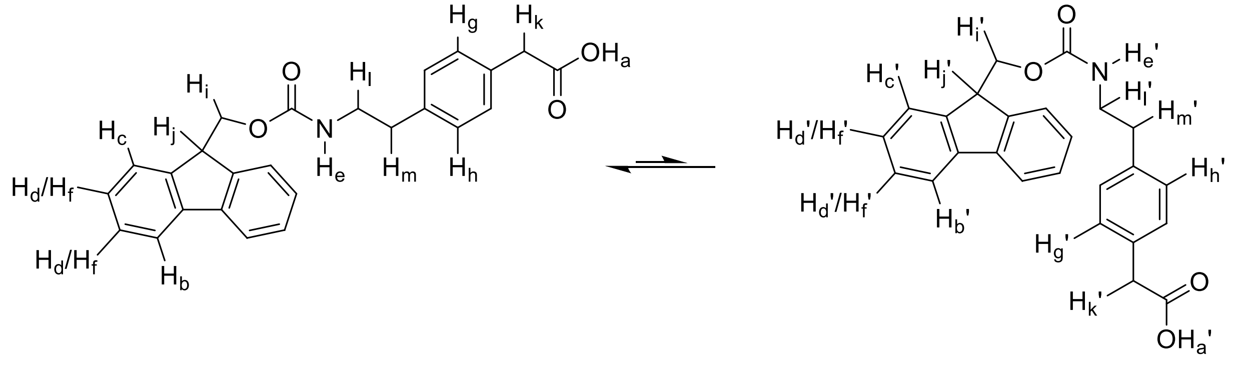

**COSY correlations:**

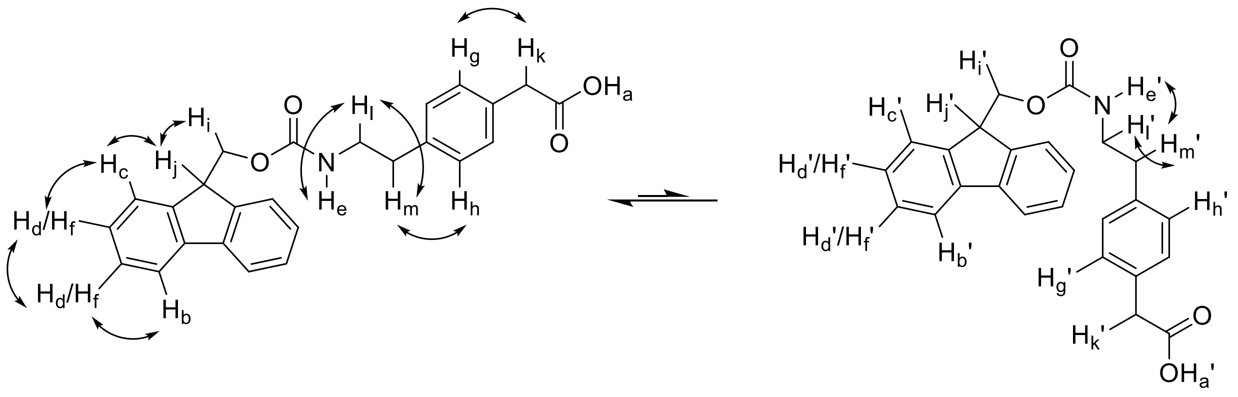

**Rotamer indicative EXSY correlations:**

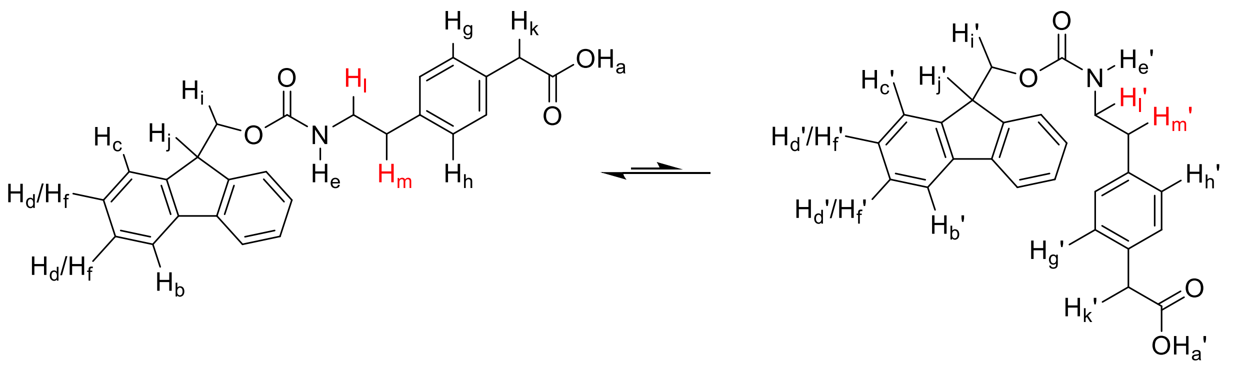

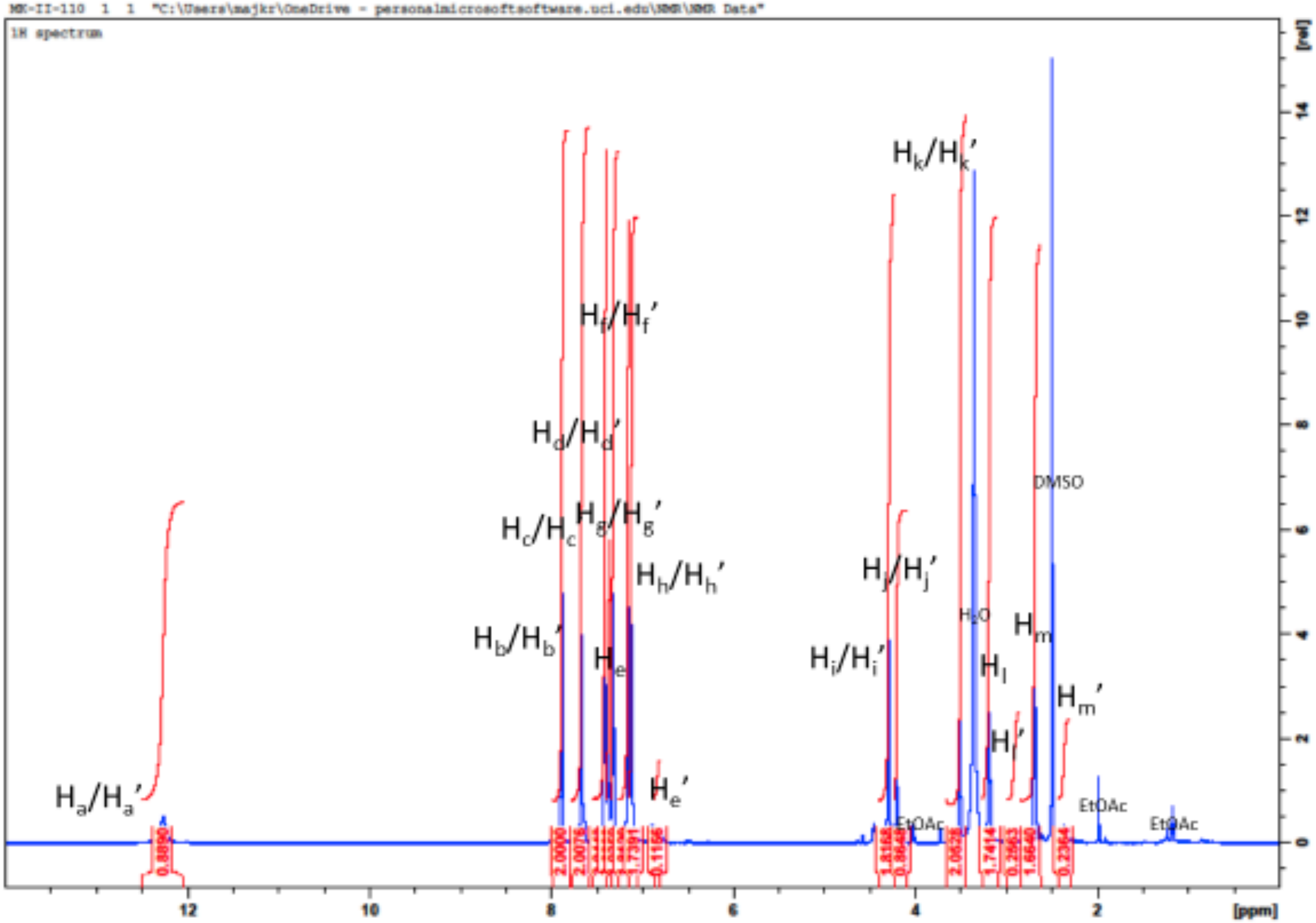

MK-II-110 1 1 "C:\Users\majkr\OneDrive - personalmicrosoftsoftware.uci.edu\NMR\NMR Data"
1H spectrum
Ha/Ha'
Hb/Hb'
Hc/Hc
Hd/Hd'
He
He'
Hf/Hf'
Hg/Hg'
Hh/Hh'
Hi/Hi'
Hj/Hj'
Hk/Hk'
Hl
Hl'
Hm
Hm'
H2O
DMSO
EtOAc
EtOAc
EtOAc
0.8890
2.0000
2.0076
1.7391
0.1166
1.8168
0.8648
2.0628
1.7414
0.2963
1.6640
0.2364
[ppm]
[rel]

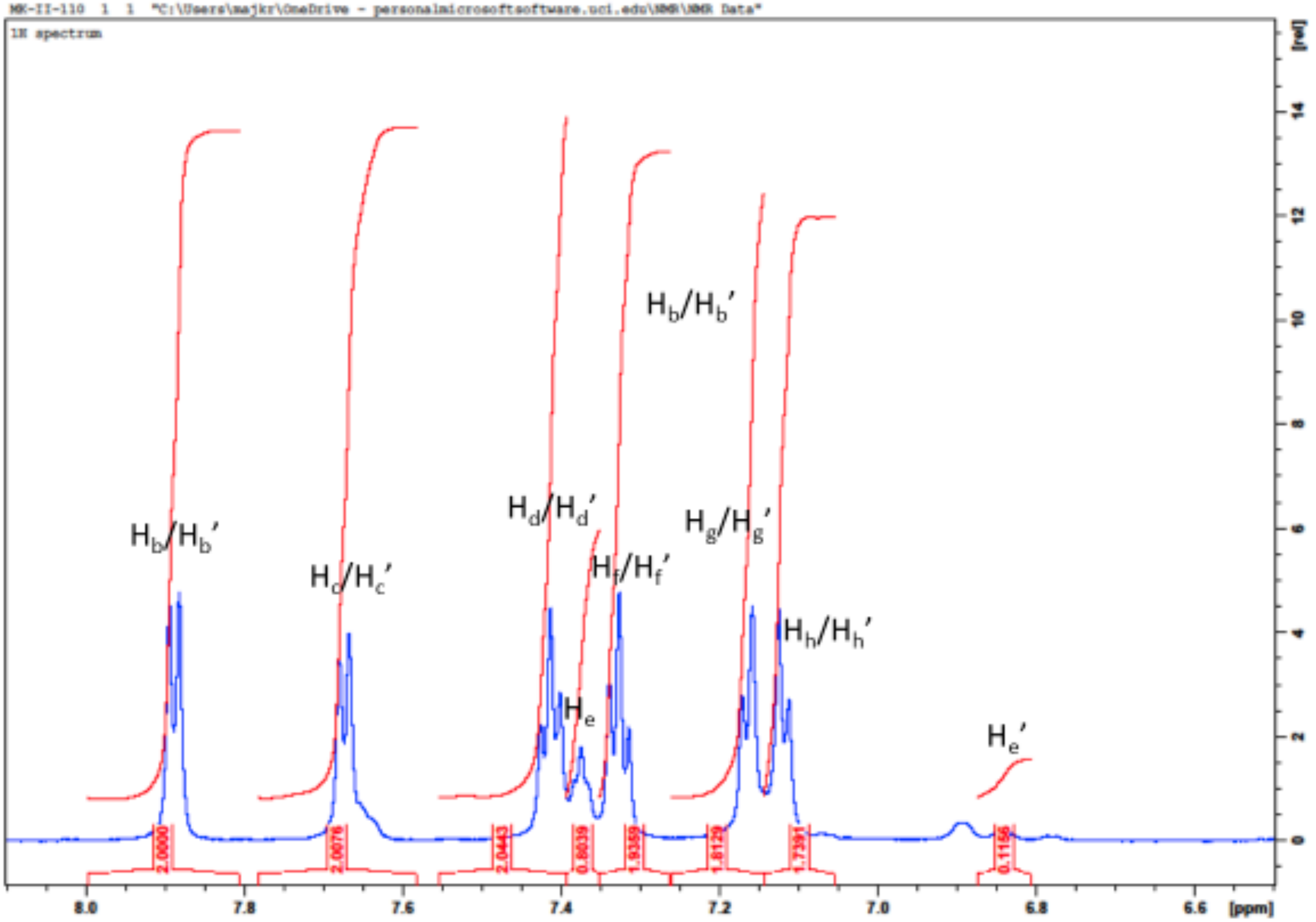
MK-II-110 1 1 "C:\Users\majkr\OneDrive - personalmicrosoftsoftware.uci.edu\NMR\NMR Data"
1H spectrum
H_b/H_b'
H_c/H_c'
H_d/H_d'
H_e
H_f/H_f'
H_b/H_b'
H_g/H_g'
H_h/H_h'
H_e'
2.0000
2.0076
2.0443
0.8039
1.9359
1.8129
1.7391
0.1156
[ppm]
[rel]

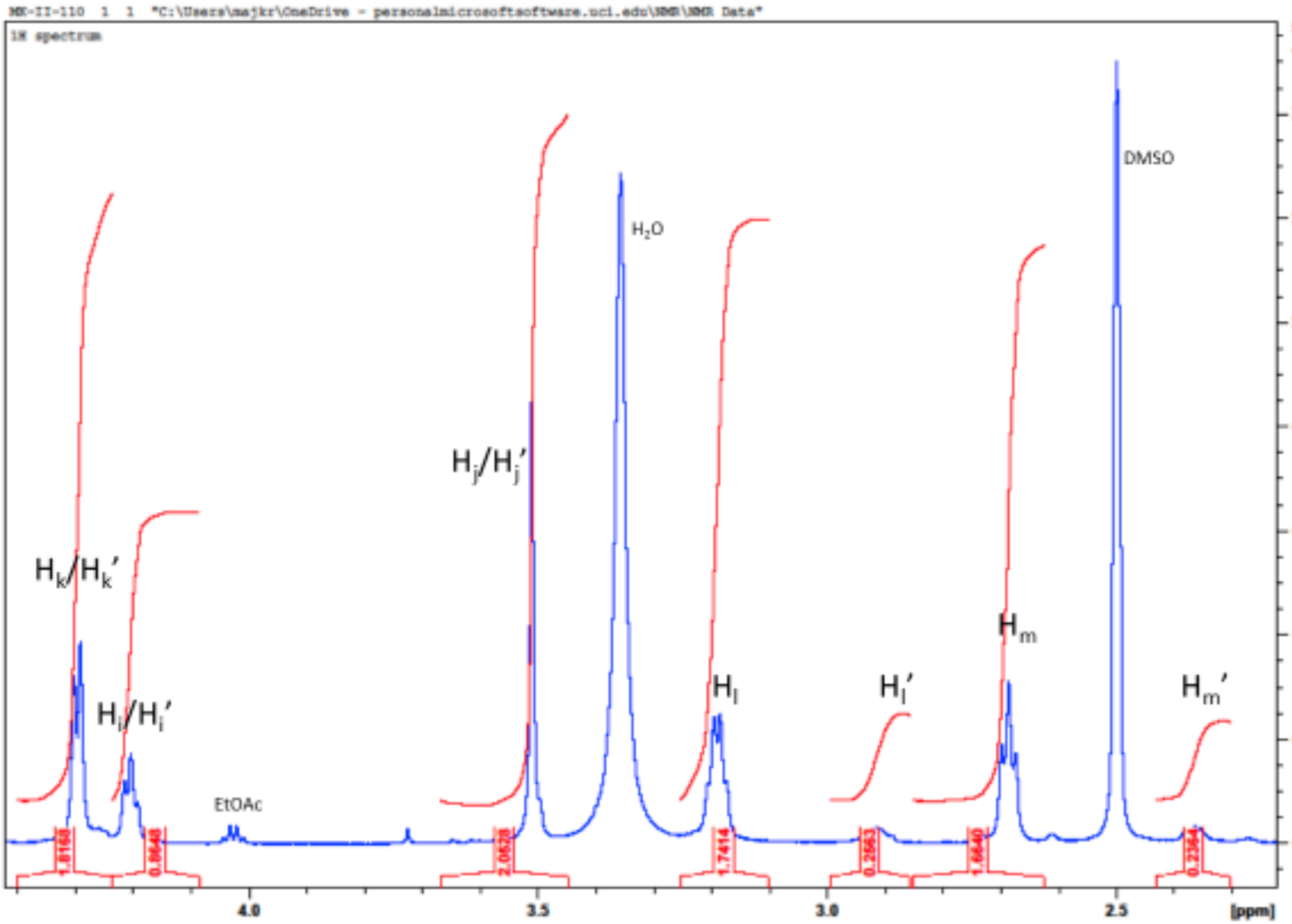
MX-II-110 1 1 "C:\Users\majkr\OneDrive - personalmicrosoftsoftware.uci.edu\NMR\NMR Data"
1H spectrum
H_k/H_k'
H_i/H_i'
EtOAc
H_j/H_j'
H_2O
H_l
H_l'
H_m
DMSO
H_m'
1.8168
0.8648
2.0628
1.7414
0.2663
1.6640
0.2364
4.0
3.5
3.0
2.5
[ppm]
[rel]
0
2
4
6
8
10
12
14

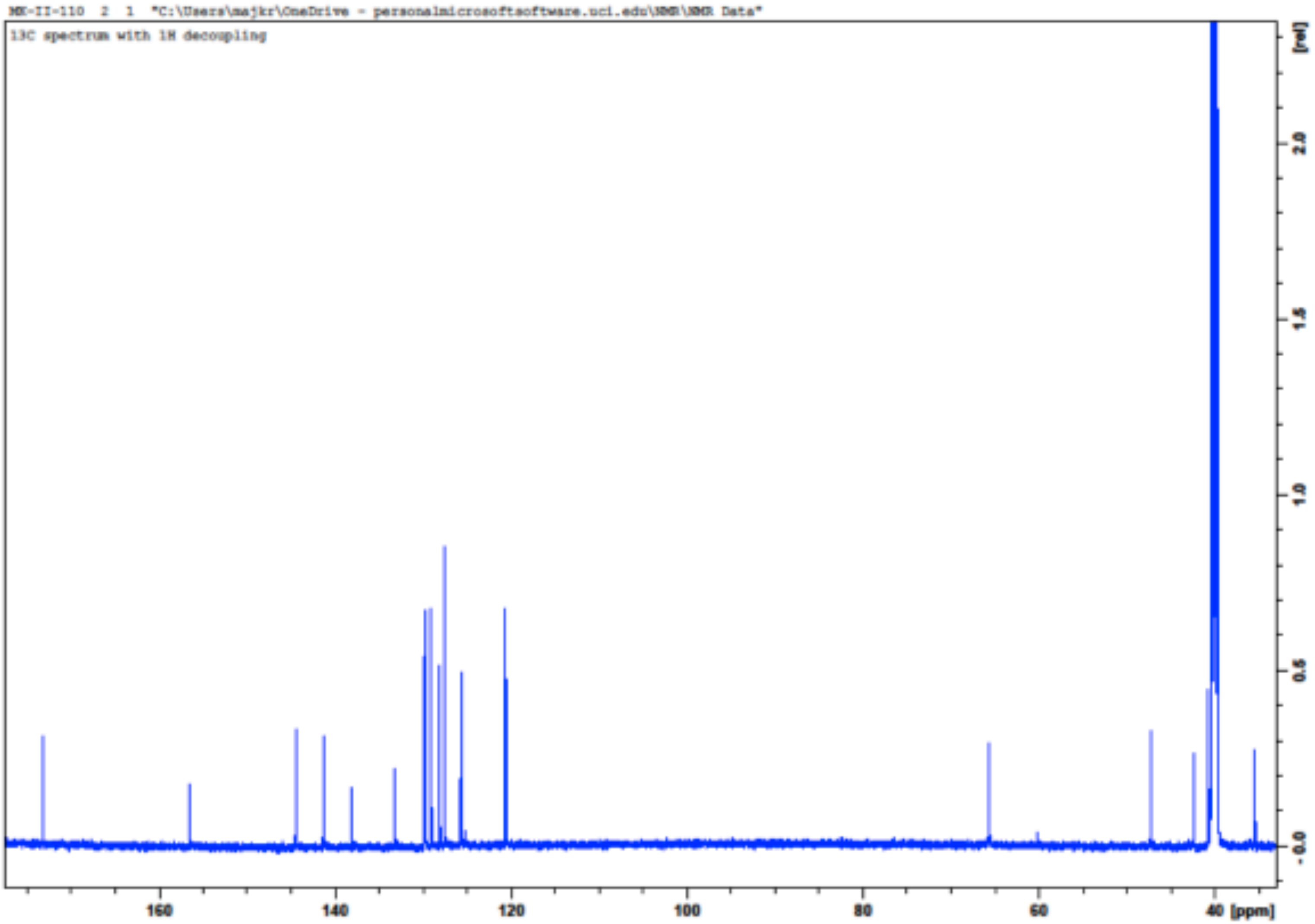
MK-II-110 2 1 "C:\Users\majkr\OneDrive - personalmicrosoftsoftware.uci.edu\NMR\NMR Data"
13C spectrum with 1H decoupling
[rel]
2.0
1.5
1.0
0.5
0.0
160
140
120
100
80
60
40 [ppm]

MK-II-110 4 1 "C:\Users\majkr\OneDrive - personalmicrosoftsoftware.uci.edu\NMR\NMR Data"

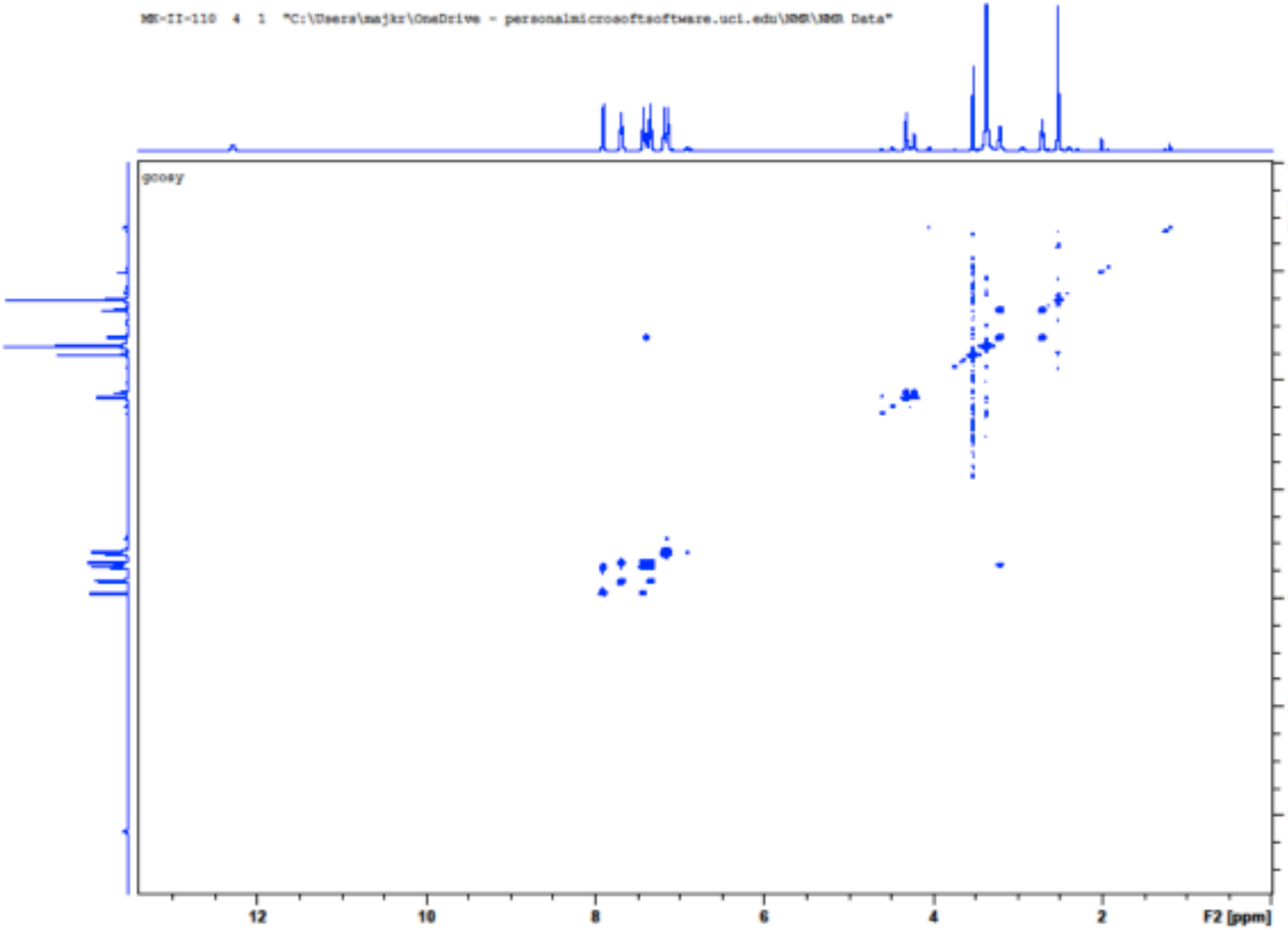

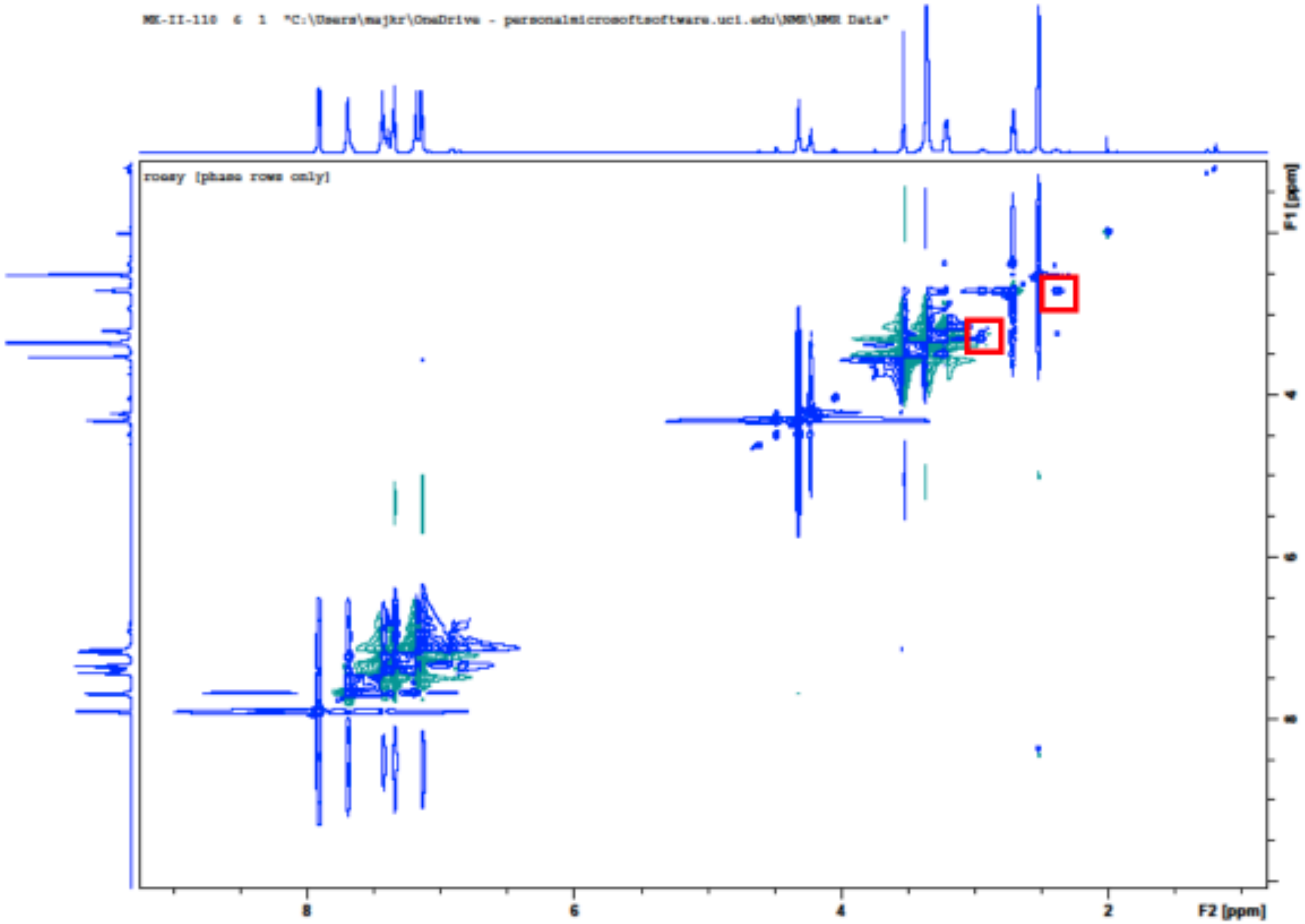
MK-II-110 6 1 "C:\Users\majkr\OneDrive - personalmicrosoftsoftware.uci.edu\NMR\NMR Data"
roesy [phase rows only]
F1 [ppm]
F2 [ppm]
8
6
4
2

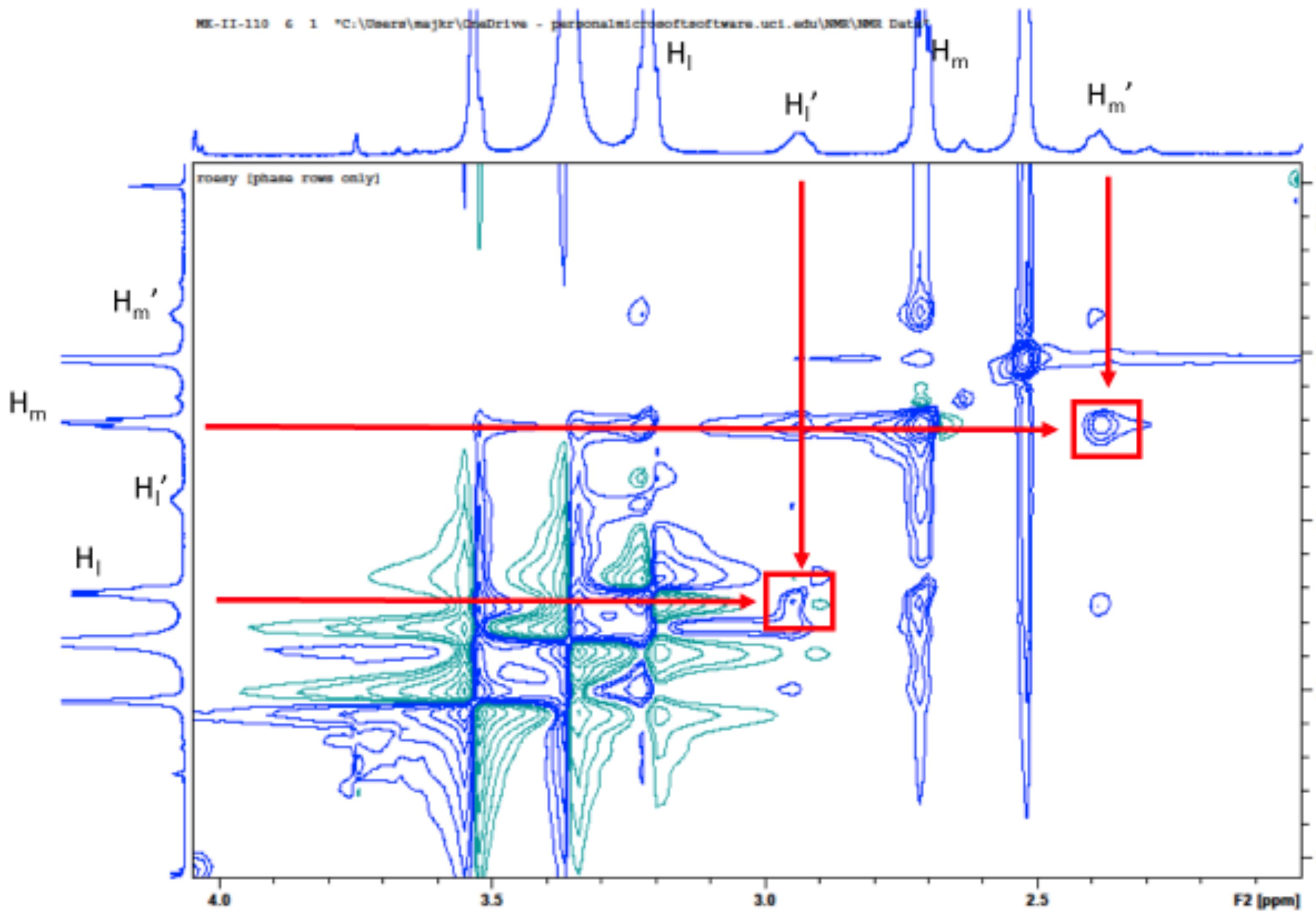
MK-II-110 6 1 "C:\Users\majkr\OneDrive - personalmicrosoftsoftware.uci.edu\NMR\NMR Data"
roesy (phase rows only)
H_l
H_l'
H_m
H_m'
F1 [ppm]
F2 [ppm]
4.0
3.5
3.0
2.5

**Peptide Characterization Data**

*Characterization of UCI-1*

Analytical HPLC trace of UCI-1.

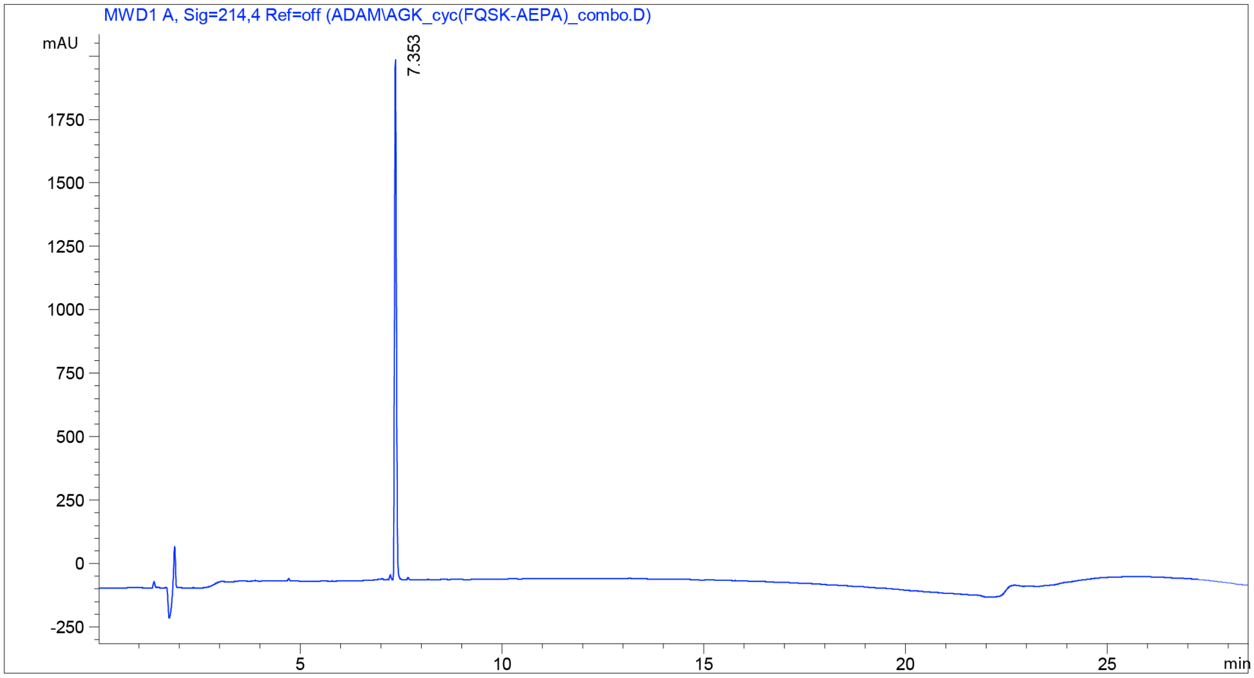

Mass spectrum of UCI-1.

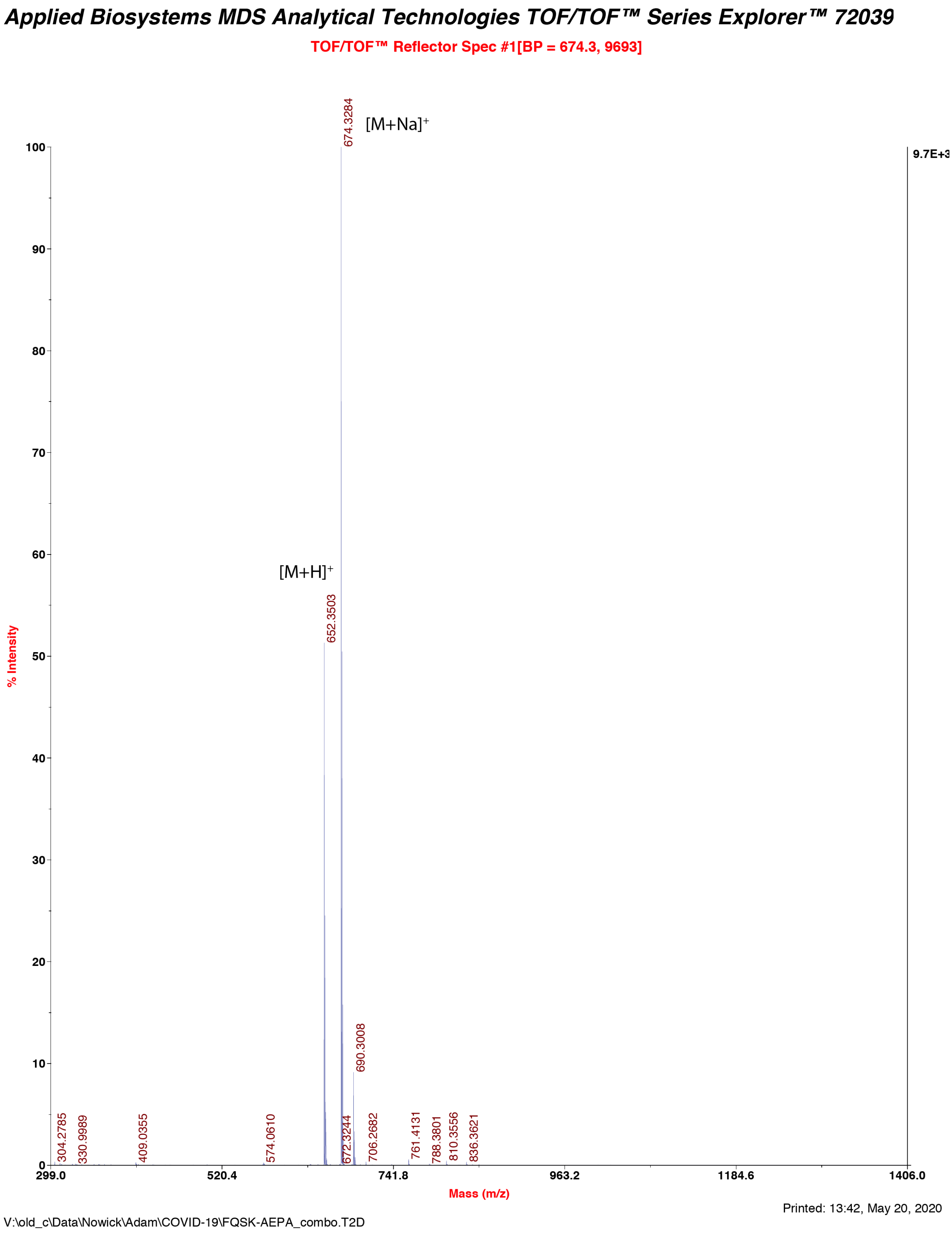

*Characterization of peptide-1a*

Analytical HPLC trace of peptide-1a.

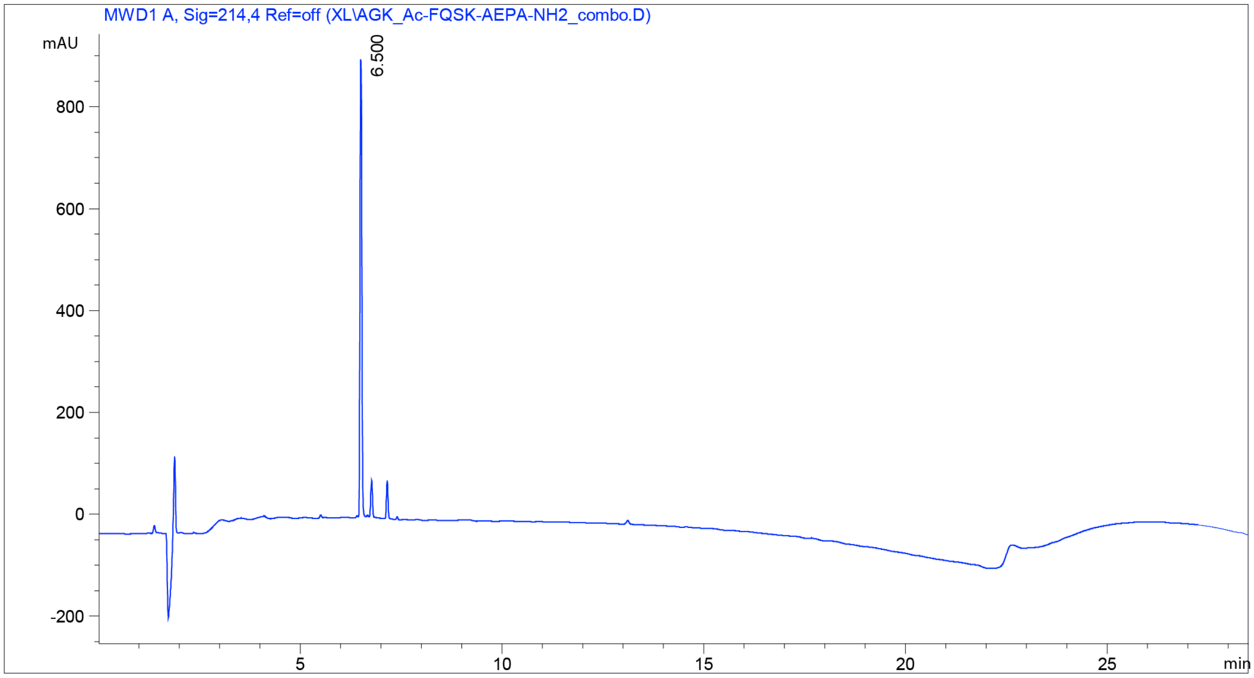

Mass spectrum of peptide-1a.

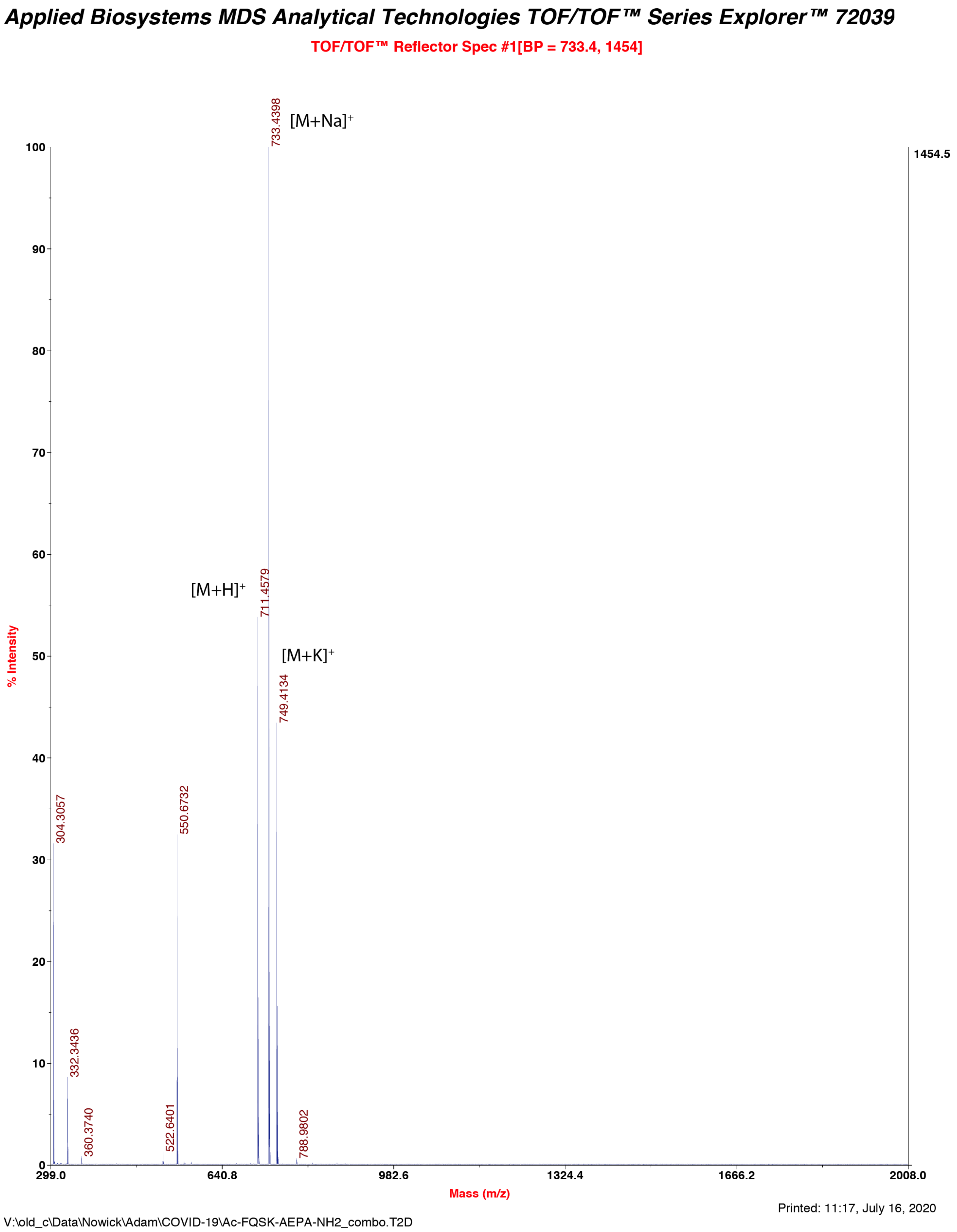

*Characterization of peptide-1b*

Analytical HPLC trace of peptide-1b.

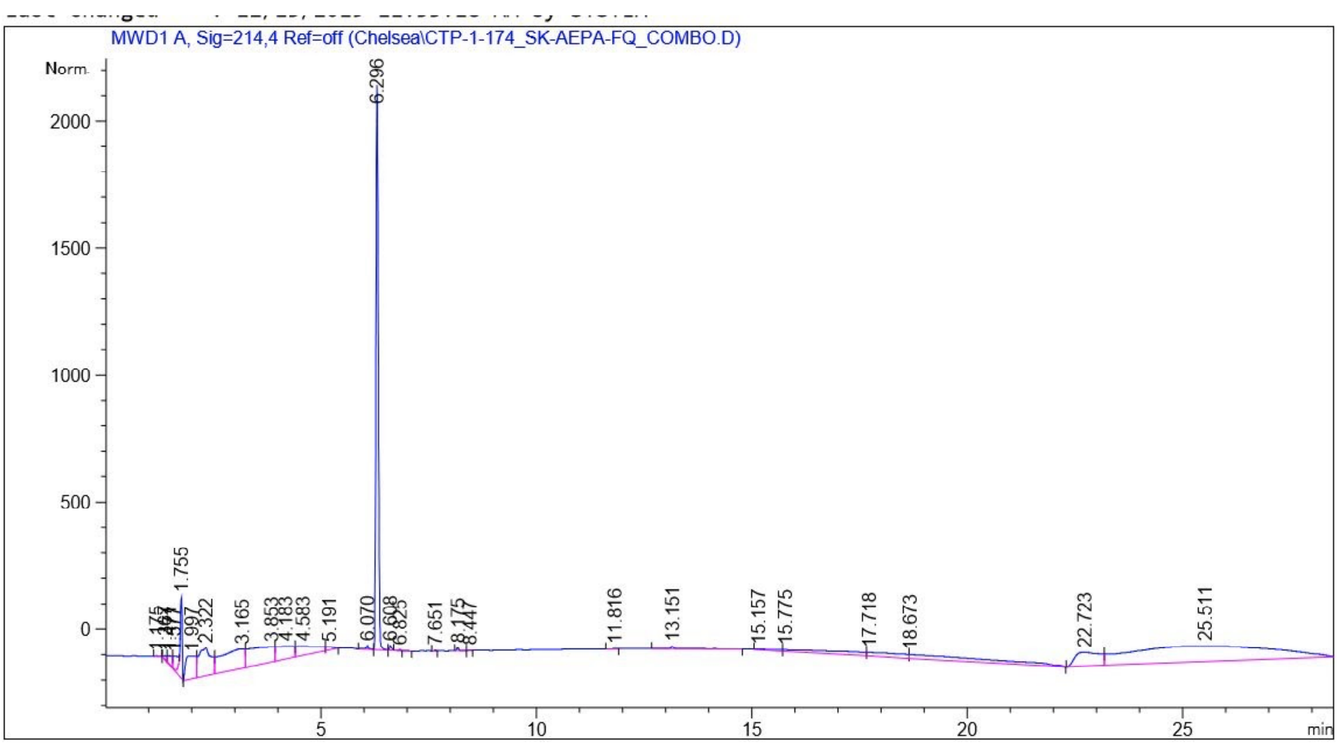

Mass spectrum of peptide-1b.

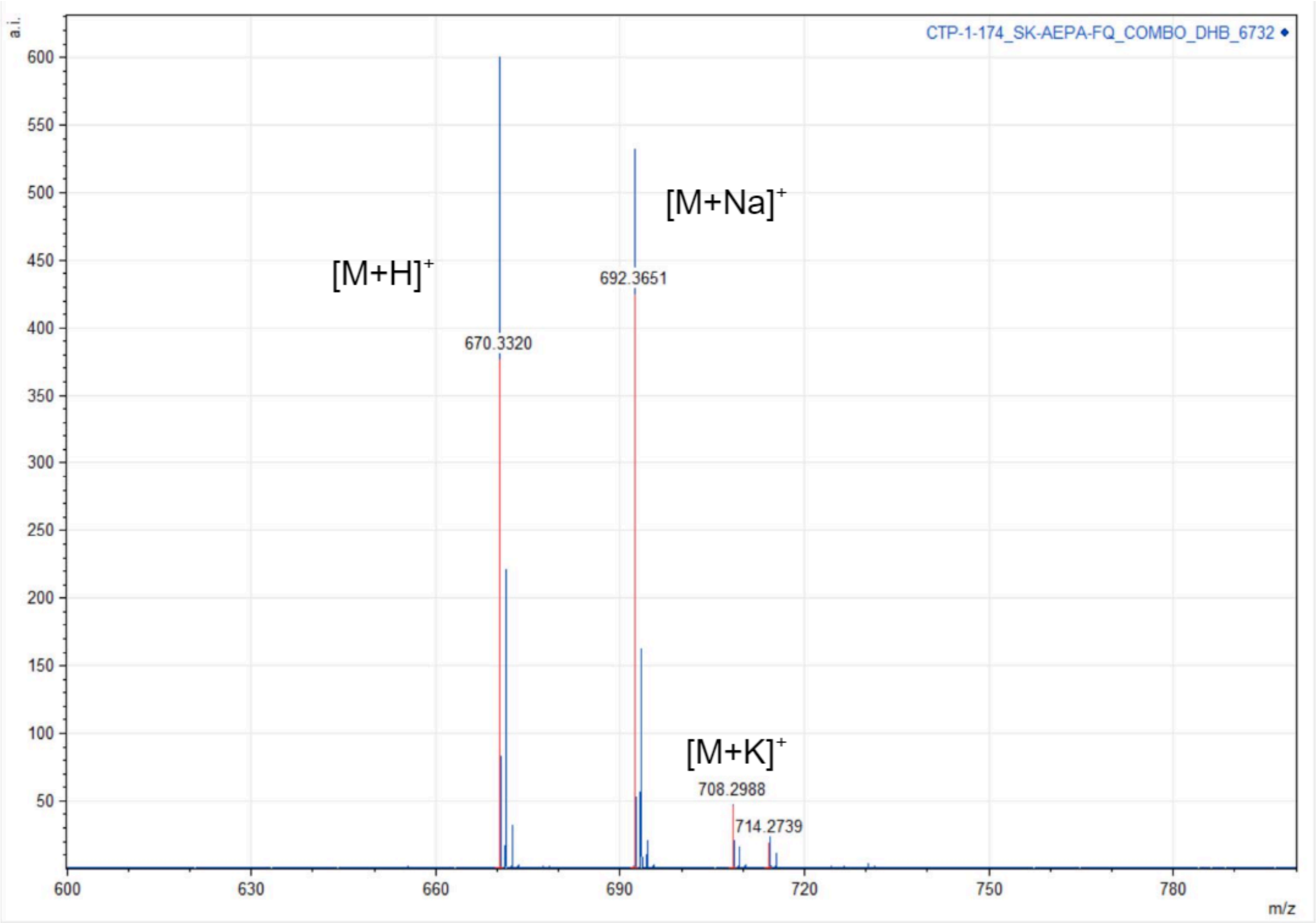
